# NanoBondy reacting through NeissLock anhydride allows covalent immune cell decoration

**DOI:** 10.1101/2025.06.14.659678

**Authors:** Lasya R. Vankayala, Kish R. Adoni, Sheryl Y.T. Lim, Omer Dushek, Konstantinos Thalassinos, Mark R. Howarth

**Affiliations:** Department of Biochemistry, University of Oxford, South Parks Road, Oxford, OX1 3QU, UK; Department of Pharmacology, University of Cambridge, Tennis Court Road, Cambridge, CB2 1PD, UK; Engineering Biology Interdisciplinary Research Centre, University of Cambridge, Cambridge, UK; Sir William Dunn School of Pathology, University of Oxford, South Parks Road, Oxford, OX1 3RE, UK; Institute of Structural and Molecular Biology, Division of Biosciences, University College London, London, WC1E 6BT, UK; Institute of Structural and Molecular Biology, Birkbeck College, University of London, London, WC1E 6BT, UK; Institute of Molecular and Cell Biology, Agency for Science, Technology and Research (A*STAR), 61 Biopolis Drive, Singapore 138673, Singapore

**Keywords:** antibody, bioconjugation, bioorganic chemistry, immunotherapy, synthetic biology

## Abstract

Cell-surface conjugation has enormous therapeutic and research potential. Existing technologies for cell-surface modification are usually reversible, non-specific, or rely on genetic editing of target cells. Here we present the NanoBondy, a nanobody modified for covalent ligation to an endogenous protein target at the cell-surface. The NanoBondy utilizes the 20 naturally occurring amino acids, harnessing NeissLock chemistry engineered from *Neisseria meningitidis*. We evaluated binding and specificity of a panel of nanobodies to CD45, a long-lived surface marker of nucleated hematopoietic cells. We demonstrated conversion of existing nanobodies to covalently reacting NanoBondies using a disulfide clamp to position the self-processing module of FrpA close to the nanobody antigen-binding site. Addition of calcium induces anhydride formation at the NanoBondy C-terminus and proximity-directed ligation to surface amines on CD45. We optimized NanoBondy reaction by fine-tuning linkers and disulfide clamp sites to modulate anhydride positioning. Tandem mass spectrometry mapped reaction sites between the NanoBondy and CD45. NanoBondy ligation was robust to buffer, pH and temperature and was detected within 2 minutes. We established reaction specificity of NanoBondies to endogenous CD45 at the surface of NK cells and T cells. NanoBondy technology provides a modular approach for targeted, inducible and covalent cell-surface modification of immune cells.

## INTRODUCTION

Molecular recognition in living systems is dominated by networks of non-covalent contacts. However, many applications in research and biotechnology are limited by instability of such binding interactions^[1,2]^. Instability may be a challenge in response to harsh conditions or force, but is most commonly an issue for long-lasting labeling, such as to change cell behavior *in vivo*^[3,4]^. There has been particular excitement about cell-surface conjugation to enhance cell therapy, given the great success of chimeric antigen receptor (CAR)-T cells against leukemia and lymphoma^[5–7]^. However, CAR-T cells have not yet fulfilled their potential in destroying solid tumors^[5–7]^. To enhance therapeutic activity, T cells have been armed with cytokines, small molecule drugs, checkpoint inhibitors, or extracellular matrix-degrading enzymes, either directly^[8–10]^ or in nanoparticles^[11–15]^ or nanogels^[16]^.

Modular tags for covalent ligation (e.g. HaloTag^[17]^, SNAP-tag^[18]^, SpyTag/SpyCatcher^[19]^, split intein^[20]^, sortase^[21]^) have been valuable for cell-surface decoration^[22–24]^. However, the bacterial origin of most tag systems raises immunogenicity concerns^[24,25]^. Genetic modification of cell therapies also faces challenges in the time from transduction to surface expression, the potential for insertional mutagenesis, and innate immune activation by nucleic acid delivery^[26]^. Each genetic change to cells adds to the delay, complexity and cost of this exceptionally expensive therapeutic class^[27]^. Cells may also be modified by inserting hydrophobic moieties into the plasma membrane, which is widely applicable but lacks specificity of insertion site or cell-type^[28,29]^. while hydrophobic tails gradually de-insert into the medium or neighboring cells^[30]^. Covalent ligation has been achieved through metabolic labeling using amino acids or carbohydrates with bio-orthogonal groups, which leads to surface display for click reaction^[13,31–33]^. Chemical crosslinkers^[11,14,15]^ or N-hydroxysuccinimide-based biotinylation followed by streptavidin labeling also allow stable cell decoration^[34]^. However such approaches modify multiple proteins, which may interfere with cell function^[4,35]^.

Nanobodies, also known as Variable Heavy domain of Heavy chain (VHH) or single domain antibodies (sdAb), are a binding scaffold typically derived from immunizing llamas, alpacas or camels. Nanobodies are an excellent platform for molecular engineering, given their small size, stability, high affinity, and ease of production in *Escherichia coli*^[36,37]^. Covalently-reactive nanobodies have been produced through unnatural amino acid mutagenesis for Sulfur Fluoride Exchange (SuFEx) reaction^[31,32]^ but face issues in scalability towards large-scale protein production^[38]^.

To create a new modality for covalent recognition of unmodified proteins, here we describe the NanoBondy, a covalently-reactive nanobody harnessing our group’s NeissLock chemistry^[39]^. NeissLock is engineered from the self-processing module (SPM) of FrpA of *Neisseria meningitidis*. Addition of calcium activates SPM, resulting in rapid autoproteolysis at an aspartate-proline bond. This step leads to formation of a highly electrophilic anhydride^[39,40]^, which then can undergo attack by water or by nucleophiles such as amines on proteins nearby (Figure 1A). NeissLock has previously been used to lock together pre-existing protein-protein interactions that are naturally present in a specific organism, where there is a high resolution structure of the complex in the Protein Data Bank (PDB)^[39]^. Here we advance NeissLock technology beyond endogenously occurring protein-protein pairs, to artificial complexes where there is no experimental structure. By engineering fusion of this SPM to a pre-existing nanobody, with precise linkers and a disulfide clamp, we enable covalent conjugation of a nanobody to its protein target in an inducible and targeted manner, after which the SPM moiety can diffuse away (Figure 1B). We optimize NanoBondy conjugation in the context of the cell-surface target CD45, which is a long-lived and broadly-expressed immune marker^[41]^. We determine key features of NanoBondy design for anhydride positioning and robustness to reaction conditions for coupling to the isolated glycosylated extracellular domain. We then validate NanoBondy coupling speed and specificity upon cell-lines and primary human immune cells. Extending the NanoBondy system, we then construct a DuoBondy, capable of covalent attachment to CD45, while including a second binder moiety allowing non-covalent attachment to the cancer checkpoint inhibitor target PD-1.

**Figure 1.**
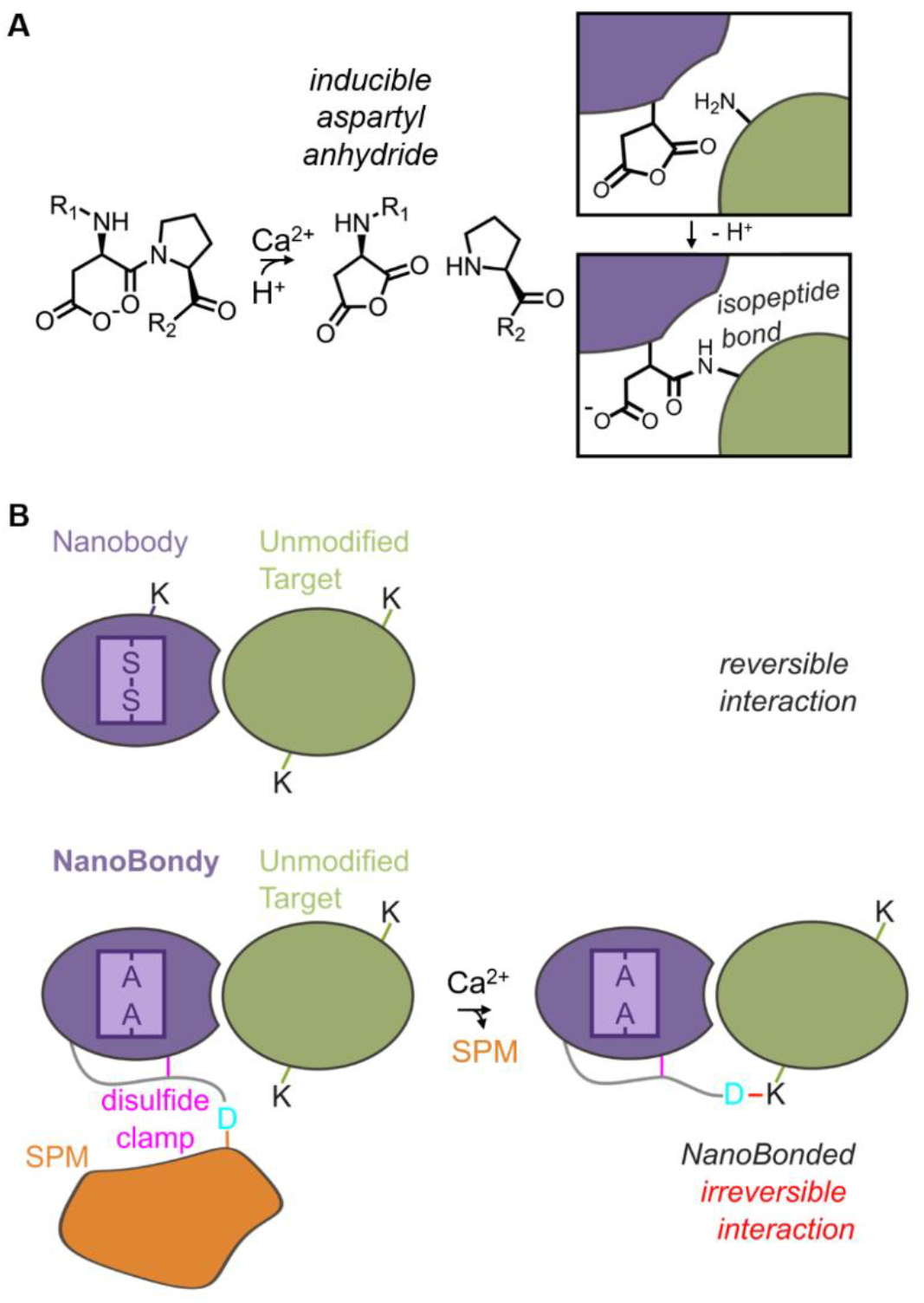
NanoBondy principle. (**A**) NeissLock chemistry. Upon addition of calcium, the self-processing module (SPM) activates autoproteolysis at the aspartate-proline bond. This step generates a highly reactive aspartyl anhydride, which undergoes nucleophilic attack by a nearby nucleophilic amino acid or water. Fusing SPM to a binder (purple) thereby allows inducible covalent coupling to a target protein (green). **(B)** NanoBondy design. A nanobody (purple) employs complementarity-determining regions (CDRs) close to the N-terminus to bind its target (green). A regular nanobody can be engineered into a covalently reacting NanoBondy by inclusion of a flexible linker and disulfide clamp (magenta) to hold the SPM (orange) near the target, allowing anhydride-mediated covalent conjugation to the target after calcium-activation.

## Results

### Selected nanobody candidates demonstrate specific binding to CD45

Nanobodies to CD45 were previously generated from llama immunization^[42]^. We selected 5 nanobodies that bind the d1d2 region of CD45, furthest from the plasma membrane and conserved across isoforms of CD45^[42]^. We cloned these nanobodies for bacterial expression with a C-terminal SpyTag003^[43]^. All five nanobodies solubly expressed in *E. coli* and were efficiently purified using SpySwitch affinity chromatography^[44]^ (Figure 2A).

**Figure 2.**
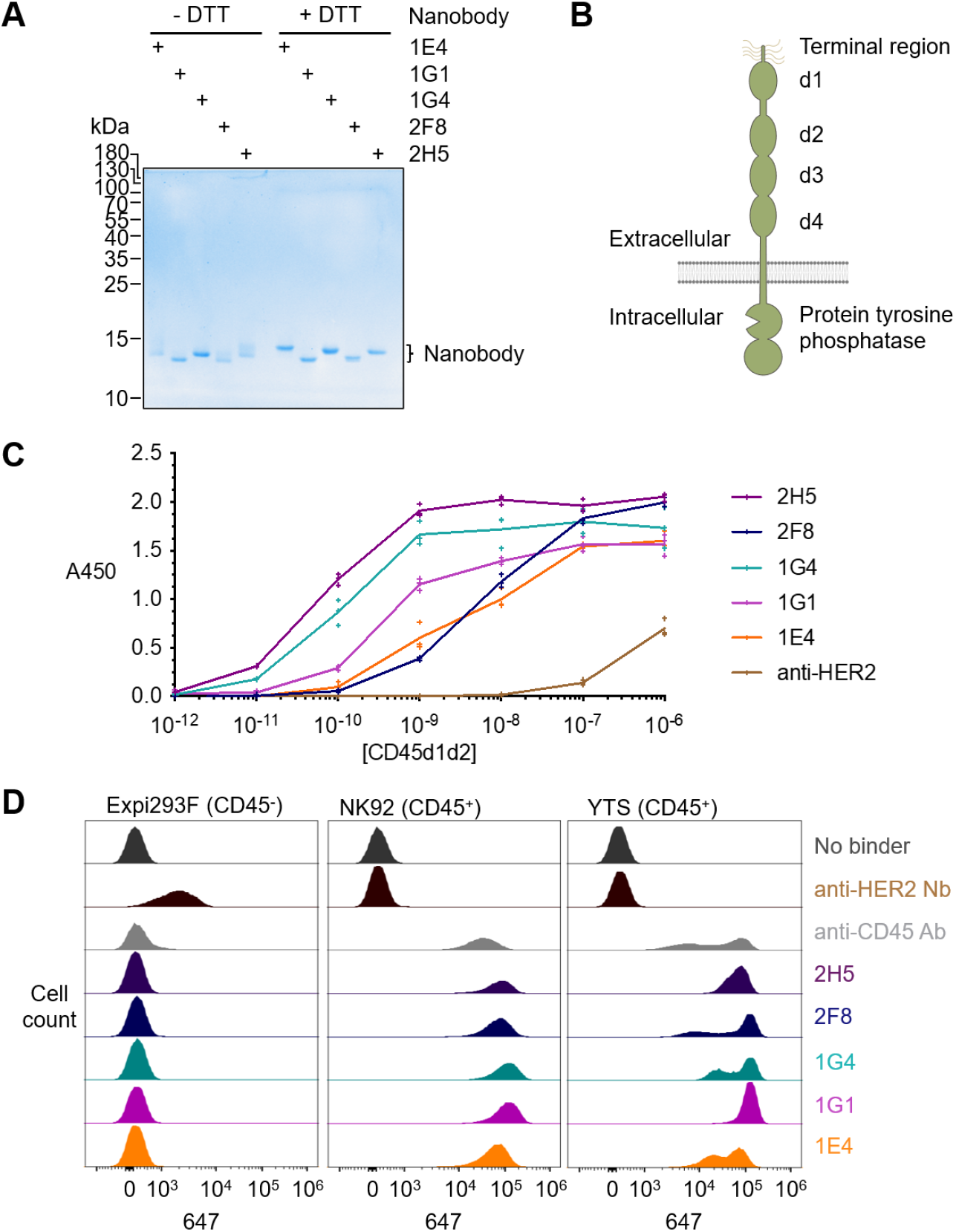
Characterization of anti-CD45 nanobodies. **(A)** Purification of anti-CD45 nanobodies. Nanobodies were expressed in E. coli and purified by SpySwitch affinity chromatography, before SDS-PAGE ± dithiothreitol (DTT) and Coomassie staining, to assess disulfide bond formation. **(B)** Schematic of the organization of CD45, including extracellular domains d1-4. **(C)** Binding of nanobodies to purified CD45. Nanobodies were coated on a plate and incubated with the indicated concentration of biotinylated human CD45 domains 1 and 2 (CD45d1d2), before colorimetric ELISA detection (absorbance at 450 nm). Anti-HER2 nanobody was a negative control. Each triplicate data point is shown, with a line connecting the mean. **(D)** Binding of anti-CD45 nanobodies at the cell-surface by flow cytometry. Anti-CD45 nanobodies were incubated with Expi293F, NK92 or YTS cells. Nanobody binding was detected using anti-VHH-Alexa Fluor 647. Anti-CD45 antibody was a positive control, with anti-HER2 nanobody or unstained (no binder) cells as negative controls.

We expressed a recombinant fragment of the d1 and d2 domains of human CD45 (Figure 2B) linked to an AviTag for site-specific biotinylation (CD45d1d2) in Expi293F cells. Nanobody binding to CD45d1d2 was evaluated by enzyme-linked immunosorbent assay (ELISA). All nanobodies demonstrated high affinity binding to CD45d1d2, with 2H5 demonstrating the best affinity (Figure 2C). The negative control anti-HER2 nanobody showed negligible background binding to CD45d1d2 (Figure 2C).

Nanobody binding to endogenous human CD45 at the cell-surface was tested by flow cytometry. Nanobodies were incubated with the YTS and NK92 natural killer (NK) cell-lines (each CD45^+^), using Expi293F cells (CD45^-^) as a negative control. All nanobodies demonstrated binding to the CD45^+^ NK cell lines, with minimal non-specific binding to CD45^-^ cells (Figure 2D). The anti-HER2 nanobody served as positive control that the HER2^+^ Expi293F cells could be stained successfully (Figure 2D). For further NanoBondy development, we prioritized 2H5 as the highest affinity binder based on ELISA, as well as high level and specific staining in flow cytometry.

### Designed NanoBondies demonstrate specific, inducible coupling to purified CD45

In the absence of experimental structures for complexes with these nanobody binders, we utilized AlphaFold2-multimer^[45,46]^ and ParaFold^[47]^ to predict docking of a 2H5-derived NanoBondy to CD45d1d2 (Figure 3A). A NanoBondy is generated by fusing FrpA SPM to the nanobody’s C-terminus via a flexible linker containing a cysteine clamp, to position the C-terminal anhydride of the NanoBondy close to the target for reaction (Figure 1B). Before the Asp-Pro cleavage site, we place a Gly-Ser-Tyr linker, which we previously established as optimal for rapid high-yielding cleavage and anhydride generation^[39]^.

**Figure 3.**
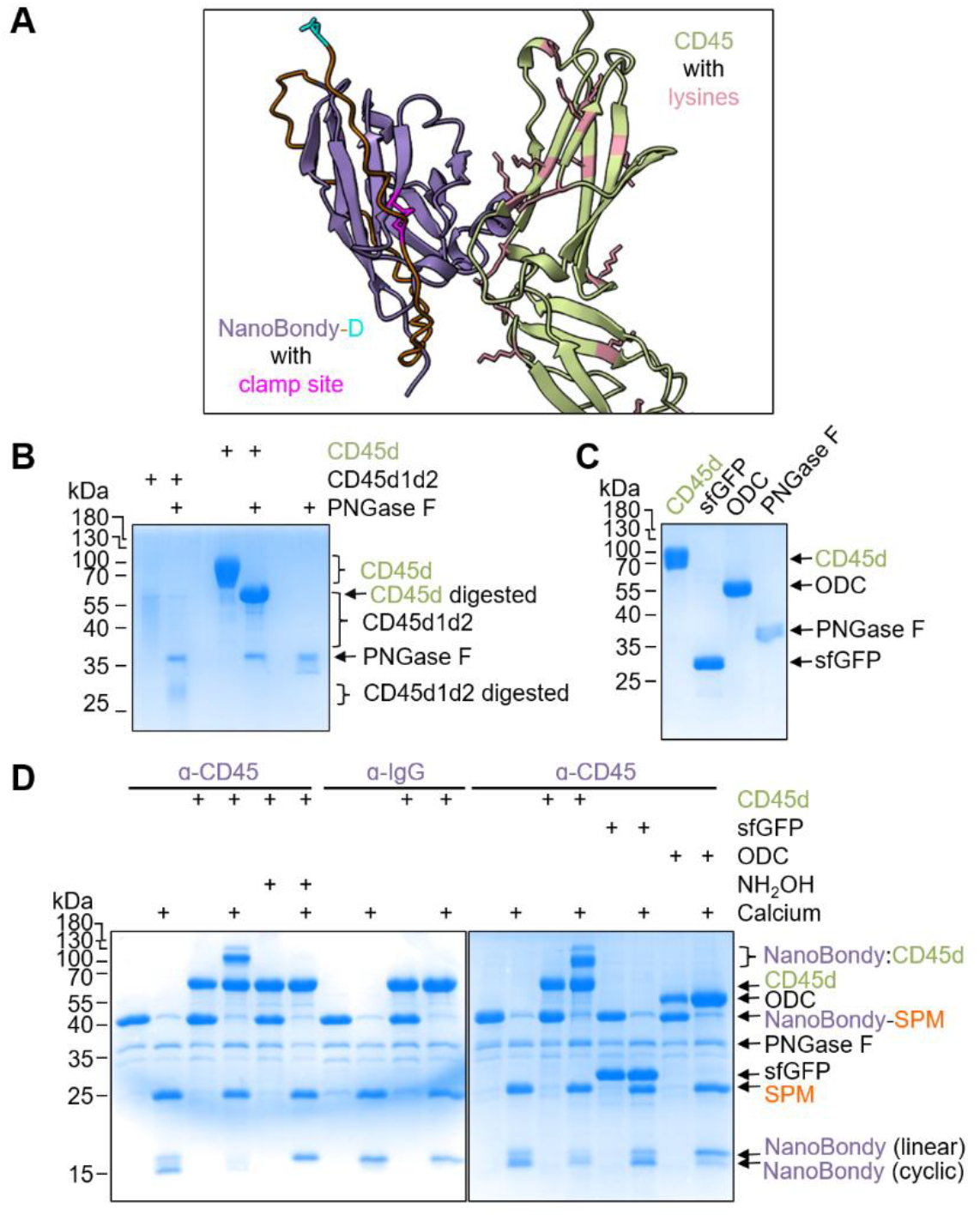
NanoBondy covalent conjugation to recombinant CD45. **(A)** AlphaFold prediction of 2H5 nanobody (purple) interaction with CD45d1d2 (green). Magenta represents the site for a disulfide clamp, with lysines on CD45d1d2 in pink and the terminal aspartate in cyan. **(B)** MBP fusion improved CD45 gel-based analysis. CD45d consists of MBP fused to domains 1 and 2 of CD45. PNGase F digestion decreased heterogeneous mobility of CD45d1d2 and CD45d upon SDS-PAGE with Coomassie staining. **(C)** Individual protein components for conjugation assay in (D). CD45d, sfGFP, ODC and PNGase F were prepared for covalent conjugation. **(D)** Specificity of NanoBondy reaction with recombinant CD45. 2H5 R72C NanoBondy-SPM was incubated with CD45d, each at 10.5 µM, for 1 h at 37 ºC in HBS ± calcium, before SDS-PAGE with Coomassie staining. ODC, sfGFP and anti-IgG NanoBondy are negative controls for reaction specificity. Hydroxylamine is a competing nucleophile to block reactivity. Colon represents a covalent conjugate.

Based on the AlphaFold model, R72C on the nanobody was identified as the initial site for the disulfide clamp. Two endogenous cysteines in the nanobody that form the core disulfide bond were mutated to alanine, to minimize any potential disulfide mispairing (Figure 1B). The designed NanoBondy (amino acid sequence in Figure S1) expressed solubly in *E. coli* and was purified using either C-tag or Ni-NTA purification. This NanoBondy was further validated by intact protein electrospray ionization mass spectrometry (Figure S2).

To allow simpler discrimination of reactant and product bands in gel assays, we cloned CD45d, which consists of CD45d1d2 with maltose-binding protein (MBP) fused at its C-terminus. CD45d expresses well in Expi293F cells with a yield of 88 mg per L of culture (Figure 3B). CD45d1d2 and CD45d both exhibit extensive glycosylation, and we showed that PNGase F digestion facilitated analysis by SDS-PAGE (Figure 3B).

To test the R72C NanoBondy reactivity to CD45d, the NanoBondy was mixed with CD45d or irrelevant protein targets in equimolar concentration, before inducing conjugation for 1 h at 37 ºC with 2 mM CaCl2, comparable in concentration to the extracellular medium^[48]^. The NanoBondy demonstrated calcium-inducible covalent conjugation to CD45d that was competed out by the strong nucleophile hydroxylamine (NH_2_OH) (Figure 3D). We have previously shown that hydroxylamine quenches the SPM-derived anhydride^[39]^. We generated a negative control NanoBondy by fusion of SPM to the anti-IgG nanobody TP1170. Covalent conjugation showed specificity for the anti-CD45 NanoBondy, with no product band when anti-IgG NanoBondy was mixed with CD45d in the presence of calcium (Figure 3D). Anti-CD45 NanoBondy did not show conjugation to the non-cognate protein targets superfolder GFP (sfGFP, expected NanoBondy covalent product mass of 44.5 kDa) or ornithine decarboxylase (ODC, expected NanoBondy covalent product mass of 70.6 kDa) (Figure 3C, D). We also showed the generality for converting different nanobodies to NanoBondies, demonstrating specificity of coupling to CD45d by a NanoBondy based on a separate anti-CD45 nanobody, 2F8 (Figure S3).

### The NanoBondy clamp site and linker length alter conjugation yield

We tested 3 alternative clamp sites in our NanoBondy design (Figure 4A), with clamp sites arranged in a tripod-like format, so that the resultant anhydride could sample various surfaces on the target. The reaction of NanoBondy with CD45d at different sites yielded different reaction bands that were resolved by SDS-PAGE. The abundance of each reaction band was altered by the choice of clamp site, with R72C demonstrating the highest reaction efficiency (Figure 4B).

**Figure 4.**
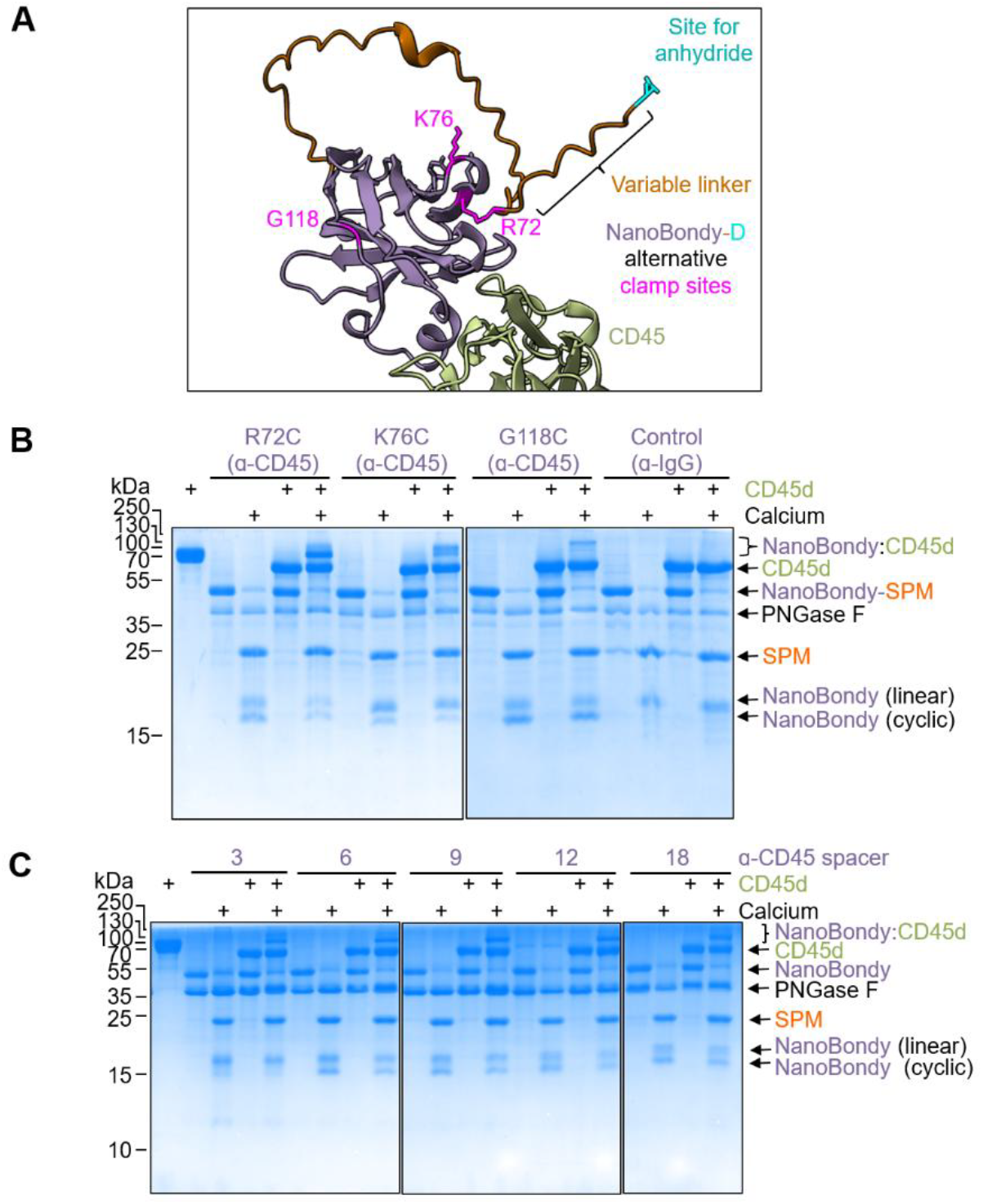
Optimization of NanoBondy clamp site and linker length. **(A)** AlphaFold prediction of 2H5 NanoBondy (purple) bound to CD45 d1d2 (green). Reactive anhydride is shown in cyan, alternative clamp sites in magenta, and linkers in orange. **(B)** Clamp-site variant reactivity. 2H5 NanoBondy variants were incubated with CD45d at 10.5 µM for 1 h at 37 ºC in HBS ± Ca^2+^, before SDS-PAGE with Coomassie staining. Anti-IgG NanoBondy is a negative control. **(C)** Linker variant reactivity for 2H5 R72C anti-CD45 NanoBondy, analyzed as in (B).

We then varied the length of the flexible linker between the cysteine clamp site and the start of the SPM, testing from 3 to 18 residues, to allow the anhydride to access more distant nucleophilic sites on the target. Interestingly, NanoBondy coupling to the CD45d target was still efficient despite these large changes in linker length (Figure 4C). From these analyses, we selected the clamp site R72C and a 9-residue linker for further exploration.

### The NanoBondy retains reactivity across various conditions

Next, we evaluated condition-dependence of NanoBondy reaction, testing covalent conjugation to its target under various situations of buffer, temperature, and pH. The NanoBondy was incubated with CD45d in the presence of calcium for varying times, before analysis by SDS-PAGE/Coomassie. The conjugation product was visible within 2 min under most conditions. Reaction proceeded more efficiently in HEPES Buffered Saline (HBS) than Tris-Buffered Saline (TBS) (Figure 5A). Reaction was faster at 37 ºC than 25 °C, with the majority of conjugation completed in 5 min at 37 °C (Figure 5A). We also evaluated pH-dependence by conducting reaction in HBS along with 2-(N-morpholino)ethanesulfonic acid (MES), to enable effective buffering over a wider pH range. NanoBondy reactivity was retained at pH 6.5-8.5, but reaction proceeded more slowly at pH 8.5 than 6.5 or 7.5 (Figure 5B). From these analyses, the optimal buffer conditions for NanoBondy reaction are HBS at 37 °C and pH 7.5. Phosphate buffered saline (PBS) is not advised for NanoBondies: addition of calcium to activate reaction would result in calcium phosphate precipitation.

**Figure 5.**
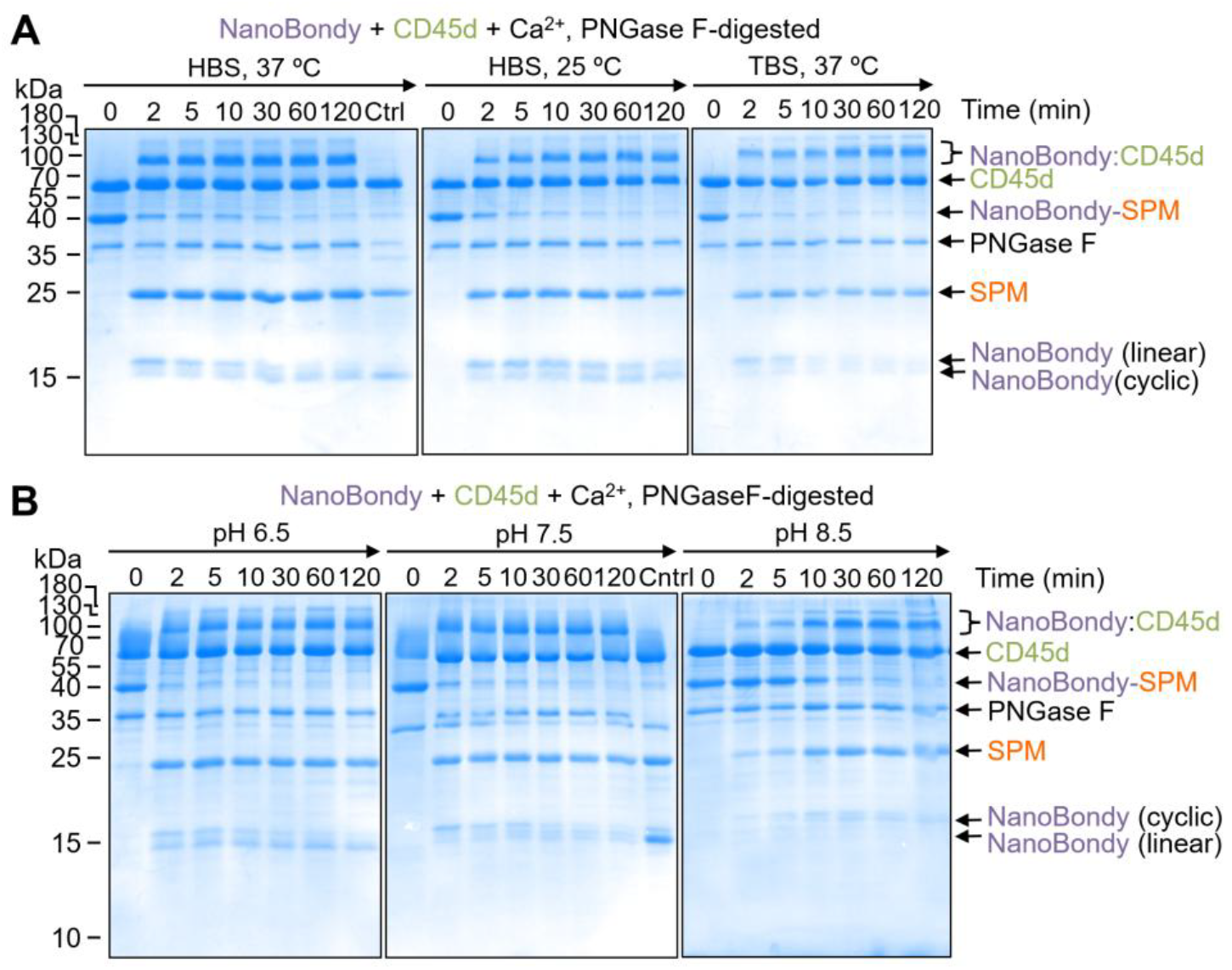
Condition-dependence of NanoBondy reaction. **(A)** Buffer- and temperature-dependence of NanoBondy reaction. 2H5 R72C anti-CD45 NanoBondy was incubated with CD45d each at 10.5 µM in the indicated buffer and temperature, before SDS-PAGE with Coomassie staining. **(B)** pH-dependence of NanoBondy reaction. Reaction was evaluated as in (A) with HBS-MES buffer at the indicated pH at 37 ºC.

### Crosslinking MS/MS identifies sites of NanoBondy-CD45d crosslinking

To identify the site of NanoBondy-CD45d crosslinking, the R72C 2H5 NanoBondy and CD45d were mixed at 10.5 µM in HBS, for a total protein content of 1 mg per reaction. The crosslinked proteins were separated by high pH reverse-phase separation and the fractions were analyzed by crosslinking tandem mass spectrometry (CL-MS/MS) to identify the dominant reaction sites (Figure 6). Crosslinking MS/MS indicated that the NanoBondy aspartyl anhydride formed covalent bonds predominantly to 4 different lysines on CD45d (Figure 6A/B, Figure S4). The crosslinked lysines were all located near the AlphaFold-predicted interface, consistent with the structure prediction (Figure 6C).

**Figure 6.**
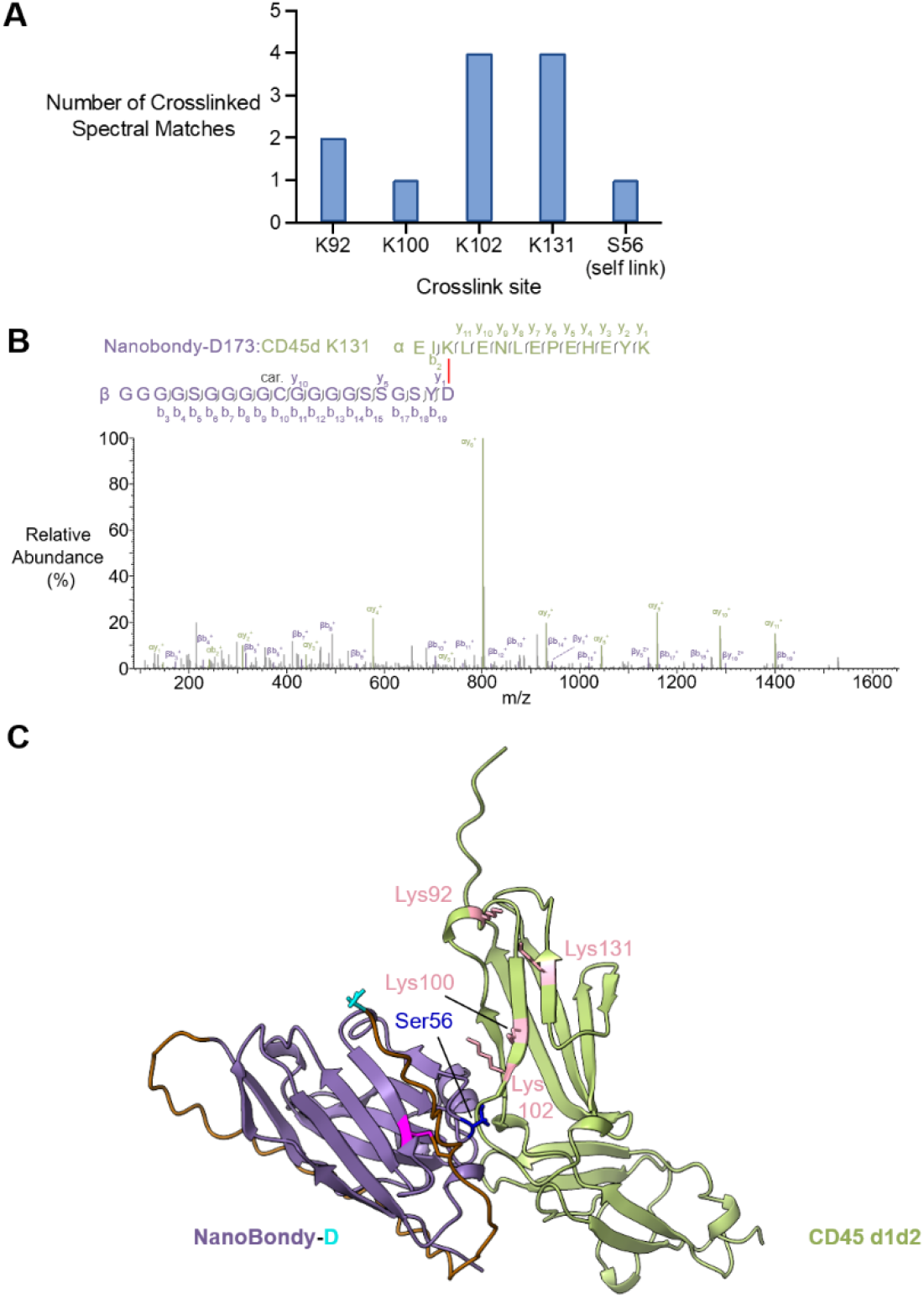
Mass spectrometry analysis of covalent conjugate between NanoBondy and CD45d. **(A)** Dominant crosslinking sites. 2H5 R72C anti-CD45 NanoBondy was incubated with CD45d, before crosslinking MS/MS. The number of identified crosslinked spectral matches for each NanoBondy crosslinking site on CD45 is shown. **(B)** Higher energy collision-induced dissociation (HCD) fragmentation spectrum of identified crosslink precursor ions corresponding to D173 (NanoBondy) to K131 (CD45d). Fragment ions matching fragmented crosslink (with crosslinker still intact) are annotated in bold, whilst peaks corresponding to fragments post-crosslink fragmentation are annotated with non-bold lines. ‘Car’ indicates carbamidomethylation of cysteine. **(C)** Mapping of crosslink sites. AlphaFold prediction of NanoBondy (purple) bound to CD45 d1d2 domains (green), highlighting crosslinking sites identified from the reactive aspartate (cyan) of the NanoBondy to target lysines (pink) or to serine (dark blue).

To further investigate these predictions, we generated point mutations of CD45d at the predicted interface with the NanoBondy (Figure S5A). The single mutants of CD45d, I104R or E105R, expressed well in Expi293F cells but cause a substantial loss in both covalent coupling by the anti-CD45 NanoBondy (Figure S5B) and non-covalent docking as tested by ELISA (Figure S5C). These mutational experiments further validated the AlphaFold-predicted interface.

Crosslinking MS/MS identified a single own-goal site, where the NanoBondy anhydride formed an ester bond with Ser56 on the NanoBondy itself (Figure 6A/B, S4D). This is the first time that NeissLock has shown covalent reaction to a serine^[39]^. We have previously shown that the SPM anhydride is reactive to nucleophiles resembling the side chains of cysteine and tyrosine, as well as to α-amines such as at the protein N-terminus^[39]^. The NanoBondy contains two cysteine residues and thirteen tyrosine residues. CD45d contains ten cysteine residues and six tyrosine residues. However, we did not observe NanoBondy-mediated conjugation to cysteine, tyrosine, or to the α-amino group on either CD45d or the NanoBondy itself.

### NanoBondy demonstrates targeted covalent coupling at the cell-surface

To test NanoBondy reaction to CD45 at the cell-surface, we initially used YTS, a human NK cell-line. Cells were incubated with NanoBondy in the presence of 2 mM CaCl_2_ for 1 h at 37 ºC. We evaluated NanoBondy reactivity to the YTS cell-surface via Western blotting. When blotting for the nanobody moiety (VHH), we consistently observed a high molecular weight band post-calcium addition, corresponding to the reaction product between NanoBondy and CD45. CD45 RO migrates at 180 kDa^[49,50]^. Therefore, the mass of the NanoBondy-D:CD45 conjugate would have an expected molecular weight of approximately 198 kDa. The observed covalent conjugate band migrated between the 185 and 270 kDa markers. We did not observe non-specific bands, indicating that the NanoBondy reaction is targeted to CD45 without promiscuous reactivity to diverse cell-surface proteins (Figure 7A).

**Figure 7.**
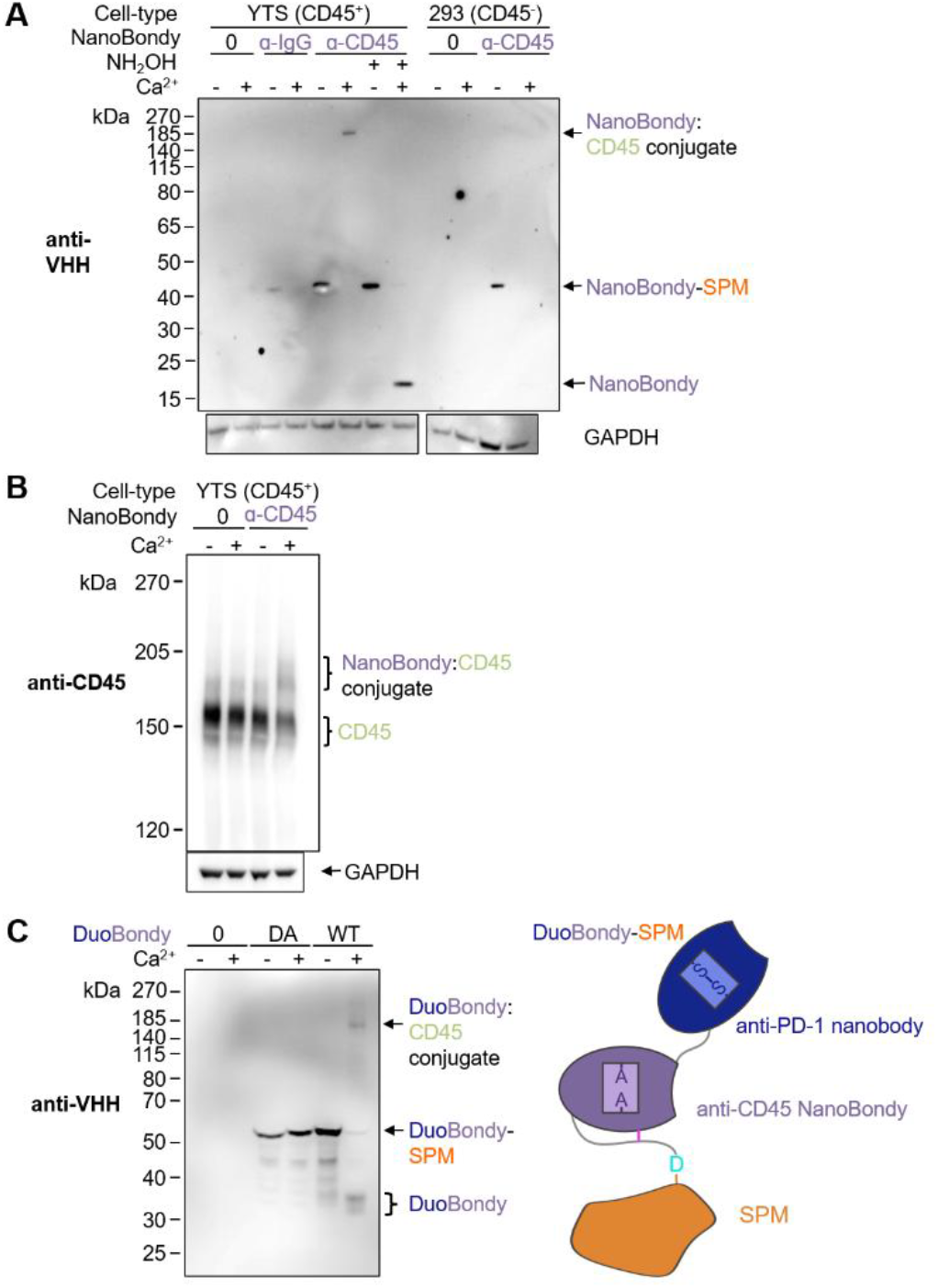
NanoBondy covalent conjugation at the cell-surface. **(A)** Western blotting of NanoBondy reaction at the surface of YTS cells. 5 µM anti-CD45 NanoBondy was incubated with YTS cells or Expi293F cells for 1 h at 37 ºC ± calcium. Covalent conjugation was evaluated by Western blot using an anti-VHH polyclonal antibody to detect the NanoBondy. Anti-IgG NanoBondy or hydroxylamine to react with the anhydride provided negative controls. Western blot to glyceraldehyde-3-phosphate dehydrogenase (GAPDH) was the loading control. **(B)** YTS cells were stained as in (A), except with 25 µM anti-CD45 NanoBondy, and detected by Western blot using an anti-CD45 antibody. The CD45 band demonstrates an upwards shift upon covalent conjugation with the a-CD45 NanoBondy. **(C)** Western blotting of DuoBondy reaction on CD8^+^ T cells. 1 µM DuoBondy (WT) or DuoBondy (DA) was incubated with CD8^+^ T cells for 40 min at 37 ºC ± calcium. Covalent conjugation was evaluated by Western blot using an anti-VHH polyclonal antibody to detect the DuoBondy. DuoBondy consists of a nanobody binder (Nb102c3) to PD-1 (dark blue) fused N-terminally to the established covalently reacting anti-CD45 NanoBondy (purple).

Conjugation was blocked upon addition of hydroxylamine, indicating that the observed product depends on anhydride-mediated reactivity. At 5 µM NanoBondy, we did not observe conjugation of the irrelevant NanoBondy control to CD45^+^ cells or conjugation of the anti-CD45 NanoBondy to the CD45^-^ Expi293F cells (Figure 7A). When blotting for CD45, we observed that the CD45 band demonstrated an upwards shift upon NanoBondy reaction (Figure 7B), supporting that the NanoBondy is reacting with endogenous CD45 at the cell-surface.

To establish NanoBondy technology for covalent delivery of effector proteins, we generated a DuoBondy, consisting of an anti-PD-1 nanobody^[51]^ attached N-terminally to the covalently reactive anti-CD45 NanoBondy (Figure 7C)^[52]^. In addition to the standard (WT) DuoBondy, we also generated a DA variant, where the reactive aspartate residue of SPM is mutated to alanine. This DA variant is therefore capable of non-covalent binding, but not calcium-mediated cleavage or conjugation. To test DuoBondy reaction, human primary CD8^+^ T cells were isolated from leukocyte blood cones. CD8^+^ T cells were incubated with 1 µM DuoBondy or DuoBondy DA in the presence of calcium. We evaluated the reactivity of the DuoBondy to the CD8^+^ cell-surface via Western blotting. The expected mass of the DuoBondy-D:CD45 conjugate is 217 kDa. When blotting for VHH, we consistently observed high molecular weight products post-calcium addition that migrated from approximately 250 kDa to just below the 185 kDa marker, consistent with the reaction product between DuoBondy and CD45 (Figure 7C). We did not observe non-specific bands, indicating that the DuoBondy reaction did not show promiscuous reactivity to other cell-surface proteins (Figure 7C). We only observed conjugation for the DuoBondy, with no covalent adduct detected for DuoBondy DA. These results demonstrate that NanoBondy technology allows covalent attachment of effector proteins to primary human cells and that fusion of an effector protein to the NanoBondy does not interfere with covalent cell conjugation.

## Discussion

Here we have established NanoBondies, re-engineering nanobodies for covalent reactivity through the inducible anhydride generation of NeissLock. We upgraded two different nanobodies to form covalent bonds to CD45. The reaction was compatible with different buffers and temperatures and showed specificity at the surface of an NK cell-line and primary human CD8^+^ T cells. Previous use of NeissLock depended on the existence of a high resolution structure in the Protein Data Bank to guide the reaction^[39]^. Despite advances in computational structure prediction, there is still uncertainty in the prediction of protein binding interfaces, particularly for contacts through antibodies and nanobodies where there are flexible loops and no evolutionary conservation^[53,54]^. This work demonstrates the harnessing of protein binders for covalent coupling where there is no experimental structure.

Binders can be upgraded to covalent reactivity by incorporation of weak electrophiles into binding interfaces, either through direct chemical coupling^[55]^ or unnatural amino acid mutagenesis^[33,56]^. Initial electrophiles such as acrylamides were highly effective for reaction to exposed cysteines at the protein-protein interface but inefficient with other side-chains^[55]^. More recent use of SuFex couples to a range of side-chains, including lysine and tyrosine^[33,57]^. However, the lack of inducible reactivity may pose challenges for sustained storage of such reagents. Similar chemical or genetic routes may also be taken to attach photocrosslinkers onto binders, but ultraviolet light activation of reactivity is damaging to cell survival and effector function^[58,59]^. NeissLock reactivity is selectively induced by gentle conditions, adding extracellular levels of Ca^2+^, and the components are fully genetically-encoded, using only canonical amino acids, which promotes simple and scalable production.

Our quantitation of cross-link frequency to different sites on the target provides valuable insights into the reach and residue-preference of the anhydride from the NanoBondy. Previous MS on anhydride reactivity from NeissLock only identified a dominant cross-link to a lysine^[39]^, but here we have quantified reaction to multiple lysine targets as well as a serine. This diverse range of possible sites for NeissLock coupling on the target is exciting, indicating that most proteins should be susceptible to ligation. Conversely, broad anhydride reactivity poses challenges in avoiding own-goal reaction sites on the NanoBondy itself when the highest reaction yield is desired. A limitation of NanoBondies is that competition between hydrolysis and coupling means that this bioconjugation is unlikely to achieve near quantitative target ligation, as may be achieved with SpyTag/SpyCatcher or HaloTag^[17,60]^.

Currently our NanoBondy optimization pipeline involves testing a small set of different clamp locations, linker lengths and own-goal lysine sites, before validating the NanoBondy with the best expression, binding specificity and reaction yield. Nanobodies or the related Sybodies are available to more than a thousand cellular targets^[61]^, so the NanoBondy strategy has potential to be generalized in diverse biological contexts. Despite internal disulfide bonds in nanobodies, we achieved efficient clamping through a novel disulfide bond in NanoBondies. In many cases the core disulfide in the nanobody was not necessary for folding and expression^[62]^ and the clamp disulfide was well formed in regular *E. coli* strains, not requiring strains optimized for an oxidizing cytosol^[63]^. Most binding scaffolds (e.g. antibodies, affibodies and DARPins)^[64]^ are like nanobodies in having the C-terminus away from the ligand binding site, so it may be feasible in future work to use clamping to engineer these other platforms into NeissLock-based covalent binders.

CD45 represents an attractive initial target for covalent cell coupling because of its high expression on a wide range of hematopoietic cells and its stable surface expression^[14]^. Future work may explore anti-CD45 NanoBondies on other cell-types, given the cancer targeting potential of CAR-macrophages and CAR-neutrophils^[65]^. Future covalent targeting will also be valuable on red blood cells, which circulate for months and lack turn-over of their plasma membrane^[35]^. Beyond cell therapy, for biomaterials^[66]^, biotransformation^[67]^ gene therapy^[68]^ and diagnostics^[69]^, target-specific irreversible coupling with NanoBondies may enable a new grade of molecular tenacity.

## ACKNOWLEDGEMENTS

L.R.V. was funded by MSD. M.R.H. was funded by the Engineering and Physical Sciences Research Council (EPSRC EP/W01565X/1). O.D. was funded by a Wellcome Trust Senior Fellowship in Basic Biomedical Sciences (207537/Z/17/Z). K.A. is funded by a Wellcome Collaborative Award in Science (209250/Z/17/Z) to K.T.. The mass spectrometer used for crosslinking was funded by a Wellcome Trust Multiuser Equipment grant (221521/Z/ 20/Z) to K.T.. S.Y.T.L. was funded by an A*STAR studentship. We thank Dr. Anthony Tumber (University of Oxford Department of Chemistry) for assistance with intact protein mass spectrometry, supported by the Biotechnology and Biological Sciences Research Council (BBSRC BB/R000344/1). We thank Robert Hedley and Vasiliki Tsioligka (Don Mason Facility of Flow Cytometry, Sir William Dunn School of Pathology, University of Oxford) for their technical expertise and assistance. We thank the flow cytometry facility staff at the University of Cambridge School of the Biological Sciences for technical expertise provided. AlphaFold2multimer docking was performed using resources provided by the Cambridge Service for Data Driven Discovery (CSD3). CSD3 is operated by the University of Cambridge Research Computing Service provided by Dell EMC and Intel using Tier-2 funding from the EPSRC (capital grant EP/T022159/1) and DiRAC funding from the Science and Technology Facilities Council. For the purpose of Open Access, the author has applied a CC BY public copyright license to any Author Accepted Manuscript (AAM) version arising from this submission.

## AUTHOR CONTRIBUTIONS

L.R.V. performed all experiments, except K.R.A. performed crosslinking MS. S.Y.T.L. developed FrpA fusion. L.R.V., O.D., K.R.A., K.T. and M.R.H. designed the project. L.R.V. and M.R.H. wrote the manuscript. All authors read and approved the manuscript.

## COMPETING INTERESTS STATEMENT

S.Y.T.L. and M.R.H. are inventors on a patent application regarding inducible anhydride formation for targeted ligation (UK Intellectual Property Office Patent Application No. 2504781.2 and 2003683.6). All other authors do not have competing interests.

