## Supplementary material for "NanoBondy reacting through NeissLock anhydride allows covalent immune cell decoration": Methods and Supplementary Figures

### Plasmids and cloning

PCR was conducted using Q5 High-Fidelity 2× Master Mix (New England Biolabs). DNA primers and gBlock gene fragments were ordered from Integrated DNA Technologies. All open reading frames were verified by Sanger sequencing (Source Bioscience and Azenta). The CD45 nanobody sequences were as described[1]. 1E4, 2H5, 2F8, 1G4 and 1G1 gBlock gene fragments were cloned into pET28a containing a C-terminal SpyTag003 (RGVPHIVMVDAYKRYK)[2] using Gibson assembly, yielding pET28a-1E4-SpyTag003 (GenBank deposition in progress), pET28a 2H5-SpyTag003 (GenBank deposition in progress), pET28a-2F8-SpyTag003 (GenBank deposition in progress), pET28a-1G1-SpyTag003 (GenBank deposition in progress) and pET28a-1G4-SpyTag003 (GenBank deposition in progress). pET28a-2H5 NanoBondy-SPM variants were generated by first cloning 2H5 C22A, C96A, R72C (GenBank deposition in progress, Addgene deposition in progress), C22A, C96A, R76C (GenBank deposition in progress) and C22A, C96A, G118C (GenBank deposition in progress) by Gibson Assembly from pET28a-2H5-SpyTag003. 2H5 mutants were then cloned, followed by a flexible linker containing SpyTag003, then the sequence for full-length FrpA SPM and a C-tag. FrpA SPM consists of amino acids 298-543 of FrpA from *Neisseria meningitidis* serotype C (UniProt: P55126). 2F8 NanoBondy (C22A, C96A, N74C) (GenBank deposition in progress) was generated by Gibson assembly from pET28a-2F8(C22A, C96A)-SpyTag003. For generation of the anti-mouse IgG NanoBondy, the nanobody TP1170[3] with C22A and C96A mutations was ordered as a gBlock and cloned into the 2H5 R72C NanoBondy plasmid with TP1170(C22A,C96A) replacing 2H5(C22A, C96A, R72C). pET28a TP1170 K98C IgG NanoBondy variant (GenBank deposition in progress) was used for future experimentation.

The 2H5 R72C NanoBondy variant was used as the parental construct for all 2H5 NanoBondy linker variants. For further purification, a 2H5 R72C NanoBondy variant bearing a C-terminal $His_6$ rather than a C-tag was also generated via Gibson assembly. To clone linker variants of the 2H5 NanoBondy, the linker following SpyTag003 (GGGGSGGGGCGGGGSSGSY) was modified via Gibson Assembly to yield the following: GGGGSGGGGCGSY (3 residue), GGGGSGGGGCGSSGSY (6 residue), GGGGSGGGGCGGGGSSGASGSY (12 residue), and GGGGSGGGGCGGGGSSGASGAGSGSGSY (18 residue).

For the anti-PD-1 DuoBondy, a gBlock gene fragment of the anti-PD-1 nanobody Nb102c3[4] was ordered from Integrated DNA Technologies, then inserted N-terminal to the NanoBondy sequence using Gibson assembly. Nb102c3 and the NanoBondy sequence were separated by a GGGGSSGSYGSY linker. The DuoBondy expression vector is pET28a containing a C-terminal $His_6$[5]. We further modified the DuoBondy sequence by replacing the loop-internal SpyTag003 with a GS linker of equivalent length. We then generated anti-PD-1 DuoBondy (DA) by mutating the reactive aspartate of the FrpA sequence to an alanine. Nanobodies, NanoBondies and DuoBondies were cloned in competent *E. coli* DH5α cells. pLEXm-hCD45d1d2 was a kind gift from Simon Davis, University of Oxford[6]. pLEXm-hCD45d1d2-AviTag was cloned by inserting the sequence for AviTag C-terminally to the coding sequence for domains 1 and 2 of human CD45, using restriction enzyme cloning. Complementary oligonucleotides encoding the AviTag sequence were annealed at equimolar concentration, then phosphorylated by T4 polynucleotide kinase (New England Biolabs). The hCD45d1d2 vector was also digested using KpnI and XhoI and was simultaneously incubated with Shrimp Alkaline Phosphatase (rSAP). Enzymes were heat-killed at 80 ºC for 20 min and digestion products were purified using a Monarch PCR and DNA cleanup kit (New England Biolabs). Insert and vector fragments were ligated using T4 DNA ligase (New England

Biolabs) for 16 h at 18 ºC. pLEXm-hCD45d (GenBank deposition in progress, Addgene deposition in progress), pLEXm-hCD45d(I104R) and pLEXm-hCD45d(E105R) were cloned by inserting the sequence for MBP C-terminal to the hCD45d1d2 (WT), hCD45d1d2(I104R) or hCD45d1d2(E105R) sequence via restriction enzyme cloning. The MBP gene was amplified via PCR, using primers to insert *KpnI* and *XhoI* sites. The MBP PCR product was purified using a Monarch PCR and DNA cleanup kit (New England Biolabs), then digested using KpnI and XhoI (New England Biolabs). The pLEXm-hCD45d1d2 vector was also digested using KpnI and XhoI and was simultaneously incubated with Shrimp Alkaline Phosphatase (rSAP). Enzymes were heat-killed at 80 ºC for 20 min. Insert and vector fragments were ligated using T4 DNA ligase (New England Biolabs) for 16 h at 18 ºC. CD45d was cloned in New England Biolab Turbo cells.

For mouse CD45d1d2 (mCD45d1d2), a gBlock gene fragment of residues 203-375 of the mouse CD45 sequence was ordered from Integrated DNA Technologies. The mCD45d1d2 gene fragment was inserted by Gibson Assembly into a pcDNA3.1 vector downstream of a tPA signal sequence with a C-terminal $His_6$ tag for purification and AviTag for site-specific biotinylation. mCD45d1d2 was cloned in New England Biolab Turbo cells.

pET28a-His6-ODC-Ctag was previously described[7] and consists of human ODC1 (UniProt P11926) with an N-terminal $His_6$ tag and a C-terminal C-tag, cloned in pET28a (GenBank MW364944, Addgene plasmid ID 163614). pET28a-SpyTag003-sfGFP was also previously described[2] (Addgene plasmid ID 133454). SpyCatcher002-MBP was previously described[8] (GenBank MF974389 and Addgene plasmid ID 102831) and consists of SpyCatcher002 fused N-terminally to MBP in pET28. pET28a-nanoHER2-SpyTag003-His6 (GenBank Accession no. PP341234, Addgene plasmid ID 216312) was previously described[8].

**Bacterial Expression**

Nanobodies, NanoBondies, sfGFP and ODC were transformed into competent *E. coli* BL21 DE3 cells (Agilent). Transformed cells were grown on LB Agar plates with 50 µg/mL kanamycin for 16 h at 37 ºC. A single colony was picked from the plate and used to inoculate 10 mL LB medium containing 50 µg/mL kanamycin. This starter culture was incubated at 37 ºC with shaking at 200 rpm for 16 h or until turbid (4 h). The starter culture was then used to inoculate 1 L of LB with 50 µg/mL kanamycin. The 1 L culture was incubated at 37 ºC with 200 rpm shaking. When $OD_{600}$ reached 0.5-0.6, expression was induced with 0.42 mM isopropyl β-D-1-thiogalactopyranoside (IPTG) for 16 h at 18 ºC with 200 rpm shaking. Bacteria were pelleted by centrifugation at 4,000 g.

**Mammalian Expression**

mCD45d1d2, CD45d1d2, CD45d and CD45d mutants were expressed in Expi293F cells (Gibco). Expi293F cells were grown in Expi293F media (Gibco) under 80% humidity and 8% (v/v) $CO_2$ at 37 ºC at 95 rpm. Transfections were conducted according to manufacturer's recommendations using the Expi293F Transfection kit (Gibco). Briefly, Expi293F cells were brought to $3.0 \times 10^6$ cells/mL. 1 µg of plasmid DNA per mL of cell culture was incubated with Expifectamine reagent for 20 min at 25 °C. This plasmid-liposome mixture was then added dropwise to the prepared Expi293F cells, and the cell culture was returned to the incubator. 18-22 h following transfection, Expifectamine Transfection Enhancers 1 and 2 were added to the cell flask. On day 5 following enhancer addition, cells were pelleted by centrifugation at 300 g, and the supernatant was collected. The supernatant was spun again at 4,000 g, then sequentially syringe-filtered through 0.45 µm and 0.22 µm filters (Starlab).

**C-tag purification**

NanoBondies and ODC bait protein were purified via C-tag affinity purification. One pellet corresponding to 1 L of bacterial culture was resuspended in lysis buffer: 20 mM Tris-HCl, 150 mM NaCl at pH 7.4, supplemented with 4 mM ethylene glycol-bis(β-aminoethyl ether)-N,N,N′,N′-tetraacetic acid (EGTA), 0.1 mg/mL lysozyme, 1 mg/mL cOmplete mini EDTA-free protease inhibitor (Sigma-Aldrich) and 1 mM phenylmethanesulfonyl fluoride (PMSF). The resuspended pellet was then sonicated on ice for 1 min at 30% amplitude, with pulses of 1 s on and 1 s off. The pellet was allowed to rest on ice for 1 min, then the sonication and rest step were repeated 3 more times. The lysate was then spun at 30,000 g and the cleared lysate was collected. An Econo-PAC Chromatography Column (Bio-Rad) containing 1 mL of Capture Select C-tag XL Affinity Matrix (Thermo Fisher Scientific) was equilibrated by 2×10 column volume (CV) washes in C-tag wash buffer (20 mM Tris-HCl, 150 mM NaCl at pH 7.4). Cleared lysate was added to the column with equilibrated resin. Lysate was incubated with resin for 1 h at 4 ºC with end-over-end rotation. The resin was then washed twice with wash buffer (20 mM Tris-HCl, 150 mM NaCl, 4 mM EGTA, pH 7.4). Protein was eluted using 6 × 1 CV washes of elution buffer: 20 mM Tris-HCl, 1 M NaCl, 50% (v/v) propylene glycol and 2 mM EGTA. The amount of protein was evaluated by $A_{280}$. Extinction coefficients were based on ExPASy ProtParam[9].

Pooled elutions were dialyzed in 3.5 kDa molecular weight cut-off Spectra/Por tubing (Spectrum Labs) overnight at 4 ºC at 1,000-fold excess against TBS (20 mM Tris-HCl, 150 mM NaCl, pH 7.4) with 100 µM EGTA. This dialysis was repeated once more for 3 h. The pooled elutions were then dialyzed for a further 3 h against a 1,000-fold excess of HBS (50 mM HEPES, 150 mM NaCl, pH 7.4) with 100 µM EGTA. Dialyzed protein was concentrated using a 10 kDa molecular weight cut-off spin concentrator (Vivaspin). Protein was stored in aliquots at -80 ºC to minimize freeze-thawing. Protein concentrations were determined using $A_{280}$. Typical yield was 1 mg per L of culture for NanoBondies.

**SpySwitch purification of SpyTag003-tagged nanobodies**

Nanobody cell pellets were prepared as above but were resuspended in 1× SpySwitch buffer (50 mM Tris-HCl pH 7.5, 300 mM NaCl). The SpySwitch purification system has been described[10]. Briefly, SpySwitch resin (0.75-1 mL per 1 L bacterial culture) was equilibrated with 2×10 CV SpySwitch buffer in an Econo-PAC Chromatography Column (Bio-Rad). Clarified cell lysate was added to the equilibrated resin and incubated for 1 h at 4 ºC with end-over-end rotation. Supernatant was allowed to flow through by gravity. The resin was washed with 2× 10 CV SpySwitch buffer. For nanobody purification, pH-dependent elution was employed. 6× 1.5 CV of SpySwitch pH elution buffer (50 mM acetic acid, 150 mM NaCl, pH 5.0) was incubated with the resin for 5 min per elution. Elutions were collected into neutralization buffer (0.3 CV of 1 M Tris-HCl pH 8.0). Proteins were dialyzed as above, into TBS. Approximate yield for nanobodies was 1-2 mg per L of culture.

**Ni-NTA purification**

CD45d1d2, CD45d (CD45d1d2-MBP), CD45d binding site mutants, sfGFP-SpyTag003 and SpyCatcher002-MBP were purified by nickel-nitrilotriacetic acid (Ni-NTA) affinity chromatography. Mammalian cell supernatant was collected and prepared as outlined above. The supernatants were supplemented with 10× Ni-NTA buffer (500 mM Tris-HCl, 3 M NaCl, pH 7.8) at 10% (v/v). 1 mL of Ni-NTA agarose (Qiagen) was packed in an Econo-PAC Chromatography Column (Bio-Rad). The packed resin was equilibrated with 2 × 10 CV of Ni-NTA buffer (50 mM Tris-HCl, 300 mM NaCl, pH 7.8). Prepared mammalian supernatant was then added to the equilibrated resin and incubated for 1 h at 4 ºC with end-over-end rotation. Supernatant was allowed to flow through by gravity. The resin was then washed with 2× 10 CV of Ni-NTA wash buffer (10 mM imidazole in Ni-NTA buffer). Protein was eluted by 6× 1

CV of elution buffer (200 mM imidazole in Ni-NTA buffer). Protein was dialyzed as described for NanoBondies, twice with TBS and once with HBS. Approximate yield for CD45d is 88 mg per L culture. Approximate yield for hCD45d1d2 is 13 mg per L culture. hCD45d1d2 was biotinylated as described[11].

NanoBondy-$His_6$ and DuoBondies were purified using cOmplete His-Tag purification resin (Roche). Bacterial cell pellets were lysed and prepared as above but were resuspended in sodium phosphate buffer (50 mM $NaH_2PO_4$, 300 mM NaCl, pH 8.0) supplemented with 2 mM EGTA and 5 mM imidazole. 1 mL cOmplete His-Tag agarose (Roche) was packed in an Econo-PAC Chromatography Column (Bio-Rad). The packed resin was then equilibrated with 2× 10 CV of sodium phosphate buffer. Prepared bacterial lysates were added to the equilibrated resin and incubated for 1 h at 4 ºC with end-over-end rotation. Supernatant was allowed to flow through by gravity. The resin was washed with 2× 10 CV of wash buffer (50 mM $NaH_2PO_4$, 300 mM NaCl, 10 mM imidazole, pH 8.0). Protein was eluted by 6× 1 CV elution buffer (50 mM $NaH_2PO_4$, 300 mM NaCl, 250 mM imidazole, pH 8.0). Protein was dialyzed as described above twice in TBS and once in HBS. Approximate yield for $His_6$-tagged NanoBondy was 7 mg and for DuoBondies was 1.5-4 mg per L of culture.

**Polyacrylamide gel electrophoresis**

For protein visualization, samples were mixed with 6× SDS loading buffer [234 mM Tris-HCl pH 6.8, 24% (v/v) glycerol, 120 µM bromophenol blue, 234 mM SDS] and 50 mM dithiothreitol (DTT) and heated at 99 °C for 3 min. Unless otherwise specified, SDS-PAGE was performed using 12-16% (w/v) polyacrylamide gels at 180 V in SDS-PAGE buffer [25 mM Tris-HCl, 192 mM glycine, 0.1% (w/v) SDS] in either an XCell SureLock system (Thermo Fisher) or a Mini Gel Tank (Invitrogen). Gels were washed twice with distilled water, then stained with Brilliant Blue G-250. Gels were destained in distilled water and imaged using an iBright FL1500 imaging system (Thermo Fisher), with analysis using iBright Analysis software versions 5.01 or 5.2.0 (Thermo Fisher).

**Size-exclusion Chromatography**

C-tag or His-Tag purified proteins were injected onto a pre-equilibrated Hi-Load 16/600 Superdex 200 pg column (GE Healthcare). Samples were run on an ÄKTA Pure 25 (GE Healthcare) fast protein liquid chromatography machine at 4 ºC into HBS + 100 µM EGTA, pH 7.4. Elutions were monitored by absorbance at 230 nm, 260 nm and 280 nm. Peak analysis was conducted using SDS-PAGE and desired fractions were concentrated at 4 ºC using a 10 kDa molecular weight cut-off spin concentrator (Vivaspin). Proteins were stored at -80 ºC.

**PNGase F expression and digestion**

pOPH6, a kind gift from Shaun Lott (Addgene plasmid ID 40315)[12],was transformed into competent *E. coli* BL21 DE3, as described above, and grown on LB Agar plates with 100 µg/mL carbenicillin for 16 h at 37 ºC. Bacterial growth, induction and expression were conducted as above. A bacterial pellet corresponding to 1 L of culture was resuspended in 50 mL ice-cold periplasmic lysis solution (0.5 M sucrose, 0.1 M Tris-HCl, 1 mM EDTA, pH 8.0), then pelleted again at 3,000 g for 20 min and the supernatant discarded. The pellet was then resuspended in 50 mL ice-cold MilliQ water and incubated for 10 min on ice. To the resuspended pellet was added $MgCl_2$ to a final 1 mM and the mixture was incubated on ice for a further 10 min. The bacteria were pelleted again at 3,000 g for 20 min and the supernatant retained. This supernatant was supplemented with 10× Ni-NTA buffer (500 mM Tris-HCl, 3 M NaCl, pH 7.8) at 10% (v/v). PNGase F was purified using Ni-NTA as described above. Size-exclusion chromatography was used as a secondary purification step as above. PNGase F was obtained at ~11 mg per L of culture. Purified PNGase F was aliquoted and stored at -20 ºC in HBS + 20% (v/v) glycerol.

For digestion, to the reaction mixture was added 10× glycoprotein denaturing buffer (New England Biolabs) at 10% (v/v). Reaction was heated at 100 ºC for 10 min, then centrifuged using a benchtop mini centrifuge at 25 ºC and 2,000 g for 1 min, and cooled on ice for 2 min. To this denatured reaction was added 10× glycoprotein buffer 2 (New England Biolabs) and 10% (v/v) NP-40 (New England Biolabs), both at a final 10% (v/v). MilliQ $H_2O$ was added to reach the desired final volume. Finally, 1 µg of PNGase F was added to each reaction. Digestion was conducted at 37 ºC for 2 h.

**ELISA**

Nunc MaxiSorp plates (Thermo Fisher) were coated with 80 nM SpyCatcher002-MBP in PBS (137 mM NaCl, 2.7 mM KCl, 10 mM $Na_2PO_4$ and 1.8 mM $KH_2PO_4$, pH 7.4) and incubated for 16 h at 4 ºC. Plates were washed 3 times with TBS-T [TBS + 0.1% (v/v) Tween-20], with a 5 min incubation for each wash. Plates were blocked for 1 h at 25 °C in blocking buffer: 1% (w/v) bovine serum albumin (BSA) (Sigma-Aldritch) in TBS-T. SpyTag003-fused nanobodies were added to the blocked plates at a final 80 nM in TBS and incubated for 1 h at 25 °C. Plates were washed 3 times with TBS-T for 5 min per wash. CD45d1d2-biotin was added to the plates in 10-fold serial dilutions in TBS and incubated for 1 h at 25 °C. Plates were washed 3 more times with TBS-T, incubating for 5 min per wash. Pierce High Sensitivity Streptavidin-HRP (Thermo Fisher) diluted to 0.2 µg/mL in blocking buffer was added to the plates and incubated for 1 h at 25 °C. The plates were washed 6 times with TBS-T, with 5 min per wash. Plates were incubated at 25ºC with 1-step Ultra TMB-ELISA Substrate Solution (Thermo Scientific). The reaction was stopped at the 2 min time-point by the addition of 1 M HCl.

For ELISA in Figure S5C, a Nunc MaxiSorp plate was coated with 80 nM of CD45d variants (WT, E105R or I104R) in PBS and incubated for 16 h at 4 ºC. Plates were washed 3 times with TBS-T [TBS + 0.1% (v/v) Tween-20], with a 5 min incubation for each wash. Plates were blocked for 1 h at 25 °C in blocking buffer: 1% (w/v) bovine serum albumin (BSA) (Sigma-Aldrich) in TBS-T. 2H5 R72C NanoBondy was added to the plates in 10-fold serial dilutions in TBS and incubated for 1 h at 25 °C. Plates were washed 3 more times with TBS-T, incubating 5 min per wash. MonoRab anti-VHH HRP (GenScript, A01861) diluted to 0.2 µg/mL in blocking buffer was added to the plates and incubated for 1 h at 25 °C. The plates were washed 6 times with TBS-T, with 5 min per wash. Plates were incubated at 25 ºC with 1-step Ultra TMB-ELISA Substrate Solution (Thermo Fisher). The reaction was stopped at the 2 min time-point by the addition of 1 M HCl.

Absorbance measurements were collected at 450 nm ($A_{450}$) using a FLUOstar Omega plate reader (BMG Labtech) and Omega MARS software. Binding curves were visualized by GraphPad PRISM software (version 10.1.1).

**Cell culture for immortalized cell lines**

Expi293F cells were from Thermo Fisher. YTS cells (RRID:CVCL_D324) and NK92 cells (RRID:CVCL_2142)[13] were a kind gift from Daniel Davis (Imperial College London). Expi293F cells were grown in Expi293 media (Thermo Fisher) at 37 ºC, 80% humidity and 8% (v/v) $CO_2$ with shaking according to manufacturer's guidelines. YTS cells were grown in Roswell Park Memorial Institute 1640 (RPMI-1640) supplemented with 10% (v/v) fetal bovine serum (FBS), 2 mM L-glutamine, 10 mM HEPES, 1 mM sodium pyruvate, 4.5 g/L glucose and 1.5 g/L sodium bicarbonate. NK92 cells were grown in Minimum Essential Medium Alpha (MEM-Alpha) supplemented with 10% (v/v) FBS, 10% (v/v) horse serum, 50 mM 2-mercaptoethanol, 2 mM GlutaMAX, and 200 U/mL rhIL-2 (PeproTech). CHO-K1 cells were grown in complete Dulbecco's Modified Eagle Medium (high glucose DMEM supplemented with 10% (v/v) FBS and 100 U/mL penicillin-streptomycin). YTS and NK92 cells were grown at 37 ºC and 5% (v/v) $CO_2$. CHO-K1 cells were grown at 37 ºC and 10% (v/v) $CO_2$. Cells were passaged at 70-80% confluency. Cells were sub-cultured for less than 3 months. Cells were validated as mycoplasma-negative by PCR.

**Flow Cytometry for Nanobody binding**

YTS, NK92 and Expi293F cells were washed in PBS and seeded at 1 million cells/well of a 96 well CELLSTAR V-bottom plate (Grenier). Cells were incubated with 2 µM purified nanobody or 1 µg/mL anti-human CD45 antibody (Invitrogen, clone HI30) in flow buffer [PBS + 1% (w/v) BSA] for 1 h at 25 °C. Cells were washed 3 times in PBS by centrifugation at 150 g at 4 ºC for 3 min. Cells were then incubated with secondary antibody in flow buffer for 1 h at 25 °C. For samples incubated with mouse anti-human CD45, the secondary antibody was polyclonal Alexa Fluor647-conjugated, highly cross-adsorbed goat anti-mouse IgG (Invitrogen, RRID AB_2535805), at 4 µg/mL in flow buffer at 25 ºC. For samples incubated with nanobody, the secondary antibody was iFluor647-conjugated MonoRab™ rabbit anti-camelid VHH antibody (GenScript) at 1 µg/mL in flow buffer at 25 ºC. Cells were washed 4 times at 4 ºC. Cells were then resuspended in LIVE/DEAD Fixable Aqua (Invitrogen) at 1 µL reconstituted stain per 1 million cells and incubated for 30 min at 25 °C in the dark. Cells were washed with PBS, then resuspended in PBS. To the resuspended cells was added an equal volume of 4% (w/v) paraformaldehyde in PBS, for a final concentration of 2% (w/v) for 20 min at 25 ºC. To these fixed cells was added PBS. Cells were then centrifuged, washed once in flow buffer, then resuspended in fresh flow cytometry buffer at 25 ºC. Samples were immediately analyzed on a CytoFlex LX analyzer. Compensation was conducted at the start of each analysis. Data were collected using CytExpert version 2.5 and analyzed using FlowJo version 10.1.0. Events were gated on cells, singlets, then live cells prior to binding analysis.

**Protein conjugation assays**

Reactions were carried out in HBS at 37 °C. In cases of hydroxylamine block, reactions were carried out in HBS + 15 mM hydroxylamine (Sigma-Aldrich) at 37°C. NanoBondy and CD45d were mixed in HBS at 10.5 µM each. Reaction was started by addition of 2 mM $CaCl_2$, with 2 mM EGTA pH 8.0 added to 'no calcium' controls. Reactions were incubated for 1 h, then quenched by addition of 10× glycoprotein denaturing buffer (New England Biolabs) at 10% (v/v). Reactions were PNGase F-digested as described above, then visualized by SDS-PAGE with Coomassie staining. Crosslinking between proteins at different sites leads to different polypeptide chain organizations with distinct gel mobility, as previously shown[7].

**Protein conjugation time-courses**

Unless otherwise specified, reactions were carried out in HBS pH 7.4 at 37 °C. To measure pH-dependence, HBS was supplemented with 50 mM MES to buffer across the desired pH range. To measure buffer-dependence, reactions were also carried out in TBS pH 7.4 at 37 °C. For both pH- and buffer-dependency experiments, proteins were buffer-exchanged into the desired buffer using a Vivaspin 500 10 kDa MWCO spin concentrator (Vivaspin) prior to assembling reactions.

A reaction mix was assembled by adding NanoBondy and CD45d at 10.5 µM in the desired buffer. Reaction was held at the desired temperature and initiated with 2 mM $CaCl_2$. At each timepoint, a fraction of the reaction mix was removed and quenched by addition of 4× stop buffer (60 mM EGTA and 4× glycoprotein denaturing buffer) for a final EGTA concentration of 15 mM and final protein concentrations of 10 µM. Quenched reactions were immediately heated at 100 °C for 10 min. For the 0 min timepoint, stop buffer was added first, followed by 2 mM $CaCl_2$. Reactions were PNGase F-digested as described above, then visualized by SDS-PAGE with Coomassie staining. Protein concentration post-PNGase F-digestion was 5 μM.

**Intact protein ESI Mass Spectrometry**

For intact protein MS a RapidFire 365 platform (Agilent) was used. The Rapid365 platform consisted of a jet-stream electrospray ionization source coupled to a 6550 Accurate-Mass Quadrupole Time-of-Flight (Q-TOF) (Agilent). 5 μM of 2H5 NanoBondy in HBS was diluted 1:1 with water and formic acid was added to a final concentration of 0.9% (v/v). The protein sample was aspirated under vacuum for 0.3 s and loaded onto a C4 solid-phase extraction cartridge. Washes were conducted using 0.1% (v/v) formic acid in water for 5.5 s. The sample was then eluted onto the Q-TOF detector for 5.5 s. MassHunter Qualitative Analysis B.07.00 (Agilent) was used for data analysis. Deconvolution settings were selected as follows: deconvolute (protein), maximum entropy deconvolution algorithm, mass range 10,000.00-80,000.00 Da, m/z range was limited to 600,0000–5,000,0000 m/z, mass step was 1,000.00 Da with baseline subtraction at baseline factor 3.00, proton adduct and automatic isotope width. Expected protein molecular weight was calculated using the ExPASy ProtParam tool[9], with 2 Da subtracted to account for the formation of a disulfide bond[14].

**Crosslinking MS sample preparation**

The reaction mix was assembled by adding 2H5 R72C NanoBondy to CD45d in HBS to final protein concentrations of 10.5 μM NanoBondy and 10.5 μM CD45d. The total mass of protein in the reaction was 1 mg. Reaction was initiated with 2 mM $CaCl_2$ and held at 37 ºC for 2 h. Reaction was PNGase F-digested as described above, and PNGase F enzyme was heat-inactivated by holding the reaction at 75 ºC for 10 min. Reaction mix was transferred to -20 ºC for overnight storage and transported to mass spectrometry analysis on dry ice.

An equal volume of 5% (w/v) SDS with 100 mM Tris-HCl (pH 8.5) was added to 500 μg calcium-cleaved 2H5 R72C NanoBondy:CD45d conjugate, before the sample was incubated at 99 ºC for 10 min, followed by water bath sonication (Fisherbrand, FB11203) at 37 kHz, power 100, at 25 ºC on sweep mode for a further 10 min. Ammonium bicarbonate (pH 8.5) was added to a final 50 mM, before addition of DTT to a final 8 mM, followed by incubation at 37 ºC for 30 min. Iodoacetamide was then added to a final 20 mM and incubated for 30 min in darkness at 25 ºC before adding trypsin (Trypsin Gold, Promega) at a ratio of 50:1 analyte:trypsin, followed by overnight incubation at 37 ºC with shaking. Digestion was quenched with a final trifluoroacetic acid concentration of 0.5% (v/v), before desalting using Waters Oasis HLB Solid Phase Extraction (SPE) plate. Samples were then reconstituted in Buffer A (100 μL 10 mM ammonium formate, pH 10.0) before fractionation (ACQUITY UPLC BEH C18 VanGuard Pre-column 130Å, 1.7 μm, 2.1 mm × 5 mm) using a gradient of 4% to 35% Buffer B [80% (v/v) acetonitrile, 10 mM ammonium formate, pH 10.0] over 8 min with fractions collected every 20 s. Fractions were then pooled to a total of 8 fractions and each fraction was reconstituted in 0.1% (v/v) formic acid before LC-MS/MS.

**LC-MS/MS (Crosslinking Mass Spectrometry)**

An UltiMate 3000 RSLCnano liquid chromatography system (Thermo Fisher), with 50 cm μPAC Neo HPLC analytical column and 0.075 mm × 20 mm trap cartridge (Acclaim PepMap C18 100 Å, 3 μm) was connected to an Orbitrap Eclipse Tribrid mass spectrometer (Thermo Fisher) via a SilicaTip emitter. Column temperature was set to 45 ºC, with an analytical column flow rate of 750 nL/min. Mobile phase A was 0.1% (v/v) formic acid with 3.2% (v/v) acetonitrile and mobile phase B was 0.1% (v/v) formic acid with 96.8% (v/v) acetonitrile. An elution gradient from 3% to 55% mobile phase B over 48 min was applied, with a total run time of 60 min. The Orbitrap Eclipse Tribrid mass spectrometer was externally calibrated using

Pierce FlexMix calibration solution and nano-ESI was performed using a SilicaTip emitter, connected to the LC *via* a HPLC liquid junction tee.

Spray stability and signal intensity were optimized by varying the SilicaTip electrospray to a final positive ion voltage of 2,000 V and modifying the Silica Tip positioning in the *x*, *y* and *z* dimensions. Transfer capillary temperature was set to 275 ºC, RF lens was set to 40%, precursor ion mass resolution was set to 120,000, precursor ion mass range was set to 350 -2000 *m/z* and precursor ion charge state was set from $3^+$ to $7^+$. $MS^1$ spectra were acquired with a Data-Dependent Analysis duty cycle (Top20), automatic gain control (AGC) was set to target 400,000 (100%), maximum injection time mode was set to Auto, precursor ions were isolated with a 1.4 *m/z* window using a quadrupole mass filter and monoisotopic precursor selection was set to peptide peak determination. For Orbitrap $MS^2$ acquisition using Higher energy Collision-induced Dissociation (HCD), collision energy was set to 30%, resolution was set to 30,000, mass range was set to normal, AGC was set to normal with an absolute AGC value of $5.00\times10^4$ and maximum injection time mode was set to Auto.

The samples were run in technical triplicate and RAW files were processed using Proteome Discoverer 2.5 (Thermo Fisher) with the Sequest HT[15] database searching node and the XlinkX[16] crosslink processing node. For Sequest HT, dynamic modifications included: oxidation (+15.995 Da, M) and N-terminal acetylation (+42.011 Da). Static modifications included: carbamidomethyl (+57.021 Da, C) with false discovery rate (FDR) set to 0.01 using the Target Decoy PSM Validator node. The FASTA file database contained sequences for 2H5 R72C NanoBondy, CD45d, PNGase F and 200 randomly selected decoy proteins for false discovery analysis. For crosslink identification, the FASTA file contained 2H5 R72C NanoBondy , CD45d and PNGase F. Relative to the FASTA file amino acid sequence, an aspartic acid and lysine/threonine/serine/arginine/tyrosine/α-amino sites were considered to crosslink with an overall chemical difference of O(-1)H(-1) from the peptides off the initial proteins 2H5 R72C NanoBondy and CD45d to the crosslinked entity, corresponding to a zero length crosslinker average mass of -17.0073 Da and a monoisotopic mass of -17.00274 Da. Crosslink residues were selected as: D-K, D-T, D-S, D-R and D-α-amino. XlinkX acquisition strategy was set to Noncleavable_fast and minimum signal/noise: 1.5. FDR was set to 0.05 in the XlinkX PD Validator node. Only high confidence crosslinks from D173 of 2H5 R72C NanoBondy were considered for further analysis and all identified crosslinks were manually verified from their corresponding MS/MS fragmentation spectra. The number of crosslinked spectral matches was reported by XlinkX, with the corresponding scan number used to manually annotate the MS/MS spectra from within the RAW file. All data are available on PRIDE (accession number PXD063277) and all validated crosslinks are provided in the Supplementary Information.

**Western blotting**

YTS cells were seeded at 1.5 million cells/well of a CELLSTAR 96 well V-bottom plate (Grenier). $CD8^+$ T cells were seeded in 15 mL falcon tubes at 4 million cells/condition. Cells were washed twice with HBS at 25ºC. Washed cells were resuspended with 5 μM NanoBondy in HBS with or without 5 mM hydroxylamine, or with 1 μM DuoBondy (WT) or DuoBondy (DA) in HBS + 1.5% (w/v) BSA. Reaction was initiated by addition of 2 mM $CaCl_2$, with 2 mM EGTA added to 'no calcium' controls. Cells were incubated at 37 °C for 1 h for NanoBondy assays or 37 ºC for 40 min for DuoBondy assays. Cells were then washed once in HBS at 25 ºC. For lysis, cells were resuspended in 180 μL/million cells ice-cold RIPA buffer [20 mM Tris-HCl, 150 mM NaCl, 1% (v/v) Triton X-100, 0.5% (w/v) sodium deoxycholate, 0.1% (w/v) SDS, pH 7.4] freshly supplemented with 1 mM PMSF and 1× cOmplete mini EDTA-free inhibitors (Roche). Resuspended cells were transferred to 1.5 mL microcentrifuge tubes and incubated on ice for 20 min. Lysed cells were then centrifuged at 12,000 g for 20 min at 4 °C to pellet nuclei. Lysate was mixed with 6× SDS loading buffer [234 mM Tris-HCl pH 6.8, 24% (v/v) glycerol, 120 μM bromophenol blue, 234 mM SDS] to a final volume of 1× SDS loading buffer. For anti-VHH visualization, 50 mM DTT was included, whereas for anti-CD45 visualization reducing agent was not used. Lysates were heated at 99 °C for 3 min before loading onto SDS-PAGE. For visualization of NanoBondy conjugation products, samples were loaded onto a 4-12% NuPAGE Bis-Tris gel (Invitrogen) and run in MOPS Buffer (Invitrogen) at 180 V for 1 h using a Mini Gel Tank (Invitrogen). For visualization of DuoBondy conjugation products, samples were loaded onto a 4-12% NuPAGE Bis-Tris gel (Invitrogen) and run in MES Buffer (Invitrogen) at 180 V for 1 h using an XCell SureLock Mini-Cell (Invitrogen). For visualization of CD45, samples were loaded onto a 6% Tris-Acetate gel and run in Tris-glycine buffer [25 mM Tris-HCl, 192 mM glycine, 0.1 % (w/v) SDS] at 180 V for 1 h using a Mini Gel Tank (Invitrogen). Gels were washed in MilliQ water, then soaked in 20% (v/v) ethanol in MilliQ water for 10-20 min with gentle rocking at 25 ºC. Proteins were transferred to PVDF membranes using an iBlot2 system (Invitrogen) and iBlot2 PVDF Transfer Stack (Invitrogen). Transfer was conducted at 15 V for 13 min.

For anti-VHH visualization, the blocking buffer was TBS with 5% (w/v) skimmed milk and 0.05% (v/v) Tween-20. For anti-CD45 and anti-GAPDH visualization, the blocking buffer was TBS with 3% (w/v) BSA and 0.05% (v/v) Tween-20. Membranes were blocked at 4 °C for 16 h. Blocked membranes were incubated with primary antibodies for 1 h at 25 °C; mouse anti-human CD45 (Invitrogen clone HI30) at 1 μg/mL or mouse anti-human GAPDH (antibodies.com, clone GA1R) at 2 μg/mL or AffiniPure goat anti-alpaca IgG VHH domain (Jackson ImmunoResearch, RRID AB_2810907) at 0.6 μg/mL. Membranes were washed 3 times with TBS-T [TBS + 0.05% (v/v) Tween-20], incubating 3 min per wash at 25 ºC. Membranes were then incubated with secondary stain for 1 h at 25 °C. For anti-CD45 and anti-GAPDH visualization, the secondary antibody was anti-mouse IgG peroxidase antibody (Sigma-Aldrich) at 1:2,500 in blocking buffer. For anti-VHH, the secondary antibody was rabbit anti-goat IgG HRP (Invitrogen, RRID AB_2534006) at 0.2 μg/mL dilution in blocking buffer. Membranes were then washed 3 times in TBS-T, incubating 3 min per wash at 25 ºC. Blots were visualized by SuperSignal West Pico PLUS Chemiluminescent Substrate (Thermo Fisher) using an iBright FL1500 imaging system (Thermo Fisher).

**Primary CD8$^+$ T cell isolation and transduction**

Leukocyte cones were purchased from the National Health Service's (UK) Blood and Transplantation service (NHS-BT). All cones were anonymized by the NHS-BT before purchase. This project has been approved by the Medical Sciences InterDivisonal Research Ethics Committee of the University of Oxford (R51997/RE001).

Human CD8$^+$ T cells were isolated from leukocyte cones using negative selection. Blood samples were incubated with 150 μL Rosette-Sep Human CD8$^+$ enrichment cocktail (STEMCELL Technologies) per mL of blood for 20 min at 25 ºC. The blood:enrichment cocktail mixture was diluted 2:1 with PBS and layered on top of Ficoll Paque Plus (GE) at a ratio of 0.8 Ficoll to 1.0 blood mixture. Falcon tubes containing the layered preparation were centrifuged at 1,200 g for 20 min at 25 ºC with slow acceleration and deceleration. The resulting buffy coat, containing isolated CD8$^+$ T cells, was collected. Cells were counted, then resuspended at 1 million cells/mL in complete RPMI: RPMI + 10% (v/v) FBS and 100 U/mL penicillin/streptomycin with 50 U/mL IL-2 (PeproTech) and CD3/CD28 human T-activator DynaBeads (Thermo Fisher) at a 1:1 bead to cell ratio. Isolated CD8$^+$ human T cells were cultured at 37 ºC and 5% (v/v) $CO_2$.

For preparation of lentivirus, HEK293T cells (ATCC) were seeded in 6 well plates in DMEM + 10% FBS and 100 U/mL penicillin/streptomycin so as to reach 60-80% confluence the following day. Media was removed from each well, so the final media volume was 2 mL complete DMEM/well of HEK293T cells. A lentiviral transduction mix was prepared containing 0.25 µg pRSV-Rev (Addgene plasmid ID 12253), 0.53 µg pMDLg/pRRE (Addgene plasmid ID 12251), 0.35 µg pVSV-G (Addgene plasmid ID 12259), 0.8 µg pHR-C259-PD1 (cloned by S. Bustamante Eguiguren, University of Oxford) and 5.8 µg X-tremeGene 360 Transfection Reagent (Roche). Prepared transduction mix was added dropwise to the seeded HEK-293T cells. 1 mL of fresh complete DMEM was added to each well after 16 h. Supernatant from each well was harvested 48 h after transduction and filtered through a 0.45 µm cellulose acetate filter.

$CD8^+$ T cell/bead mixture was seeded at 1 million cells/well of a 12 well plate in 1 mL of complete RPMI. To each well was added the supernatant from 1 well of HEK293-T cells. On day 2 and day 4 post-transduction, 1 mL of media was removed from each well of transduced T cells and replaced with 1 mL of fresh complete RPMI containing 200 U of IL2 (final IL2 concentration of 50 U/mL). On day 5 post-transduction, the Dynabeads were removed by magnetic separation. Transduced $CD8^+$ T cells were counted and resuspended at 1 million cells/mL in fresh complete RPMI with 50 U/mL IL2. Transduced $CD8^+$ T cells were counted and reseeded at 1 million cells/mL every other day for the duration of their use. T cells were used between 10 and 16 days after transduction, and any remaining cells were destroyed after day 16.

Expression of C259 TCR and PD-1 was validated by flow cytometry. Transduced T cells, as well as untransduced controls, were stained with anti-PD-1 phycoerythrin (PE) (clone EH12.2H7, RRID AB_2745543) at 0.25 ng/µL and PE-conjugated 9V peptide (1:1000 dilution, made in-house) in PBS-BSA (1%) for 20 min at 4ºC. Cells were washed once at 4 ºC with PBS, then resuspended in 70 µL PBS before analysis via flow cytometry using a BD Fortessa. Data were analyzed using Flow Jo version 10.1.0. Events were gated on cells and on singlets prior to analysis.

**Data Analysis and Visualization**

Data visualization for ELISA and flow cytometry activation curves was performed using GraphPad Prism 10 (GraphPad Software). Protein structures were visualized using PyMOL version 2.5.4 (Schrödinger). Figures for publication were generated using UCSF ChimeraX version 1.8rc202405230136[17]. NanoBondy structure and docking were predicted using AlphaFold with ColabFold version 1.5.2[18]. To generate AlphaFold2-multimer models of nanobody docking to the extracellular domain human CD45, an amino acid sequence corresponding to PDB accession code 5FN7 (domains 1 and 2) was used[6]. Figure 2B depicts the CD45 RO isoform, which corresponds to the most abundant CD45 isoform on YTS and activated $CD8^+$ T cells[19].

**Data Availability**

Plasmids encoding the NanoBondy and DuoBondy constructs have been deposited in the Addgene repository (https://www.addgene.org/Mark_Howarth/) and sequences of other constructs have been deposited in GenBank as described in the section "Plasmids and cloning". For Crosslinking MS, all data are available on PRIDE (ascension number PXD063277) and all validated crosslinks are provided in the Supplementary Information. Any requests for further information, or for resources and reagents should be directed to and will be fulfilled by the lead contact, M.R.H.

### anti-CD45 NanoBondy

```
MSGQVQLQESGGGLVQTGGSLTLSAVASGGTFSSYAMGWF    40
RQAPGKEREFVAAIGGSGDSTYYADSVKGRFTISCDNARN    80
SVYLQMNSLKPEDTAVYYAQADPTMFHKLYYGINPNEYDY    120
WGQGTLVTVSSGGGGSGGRGVPHIVMVDAYKRYKGGGGSG    160
GGGCGGGGSSGSYDPLALDLDGDGIETVAAKGFAGALFDH    200
RNQGIRTATGWVSADDGLLVRDLNGNGIIDNGAELFGDNT    240
KLADGSFAKHGYAALAELDSNGDNIINAADAAFQTLRVWQ    280
DLNQDGISQANELRTLEELGIQSLDLAYKDVNKNLGNGNT    320
LAQQGSYTKTDGTTAKMGDLLLAADNLHSRFKDKVELTAE    360
QAKAANLAGIGRLRDLREAAALSGDLANMLKAYSAAETKE    400
AQLALLDNLIHKWAETDGSSHHHHHH                  426
```

anti-CD45 2H5 (C22A, C96A, R72C, K76R)
Clamp site
Linker
Reactive D
SPM
$His_6$

***Supplementary Figure 1.*** **Amino acid sequence of lead anti-CD45 NanoBondy (2H5 R72C with 9 residue spacer).** Residue numbers begin at 1 from the N-terminal methionine, which is cleaved in the final protein sequence.

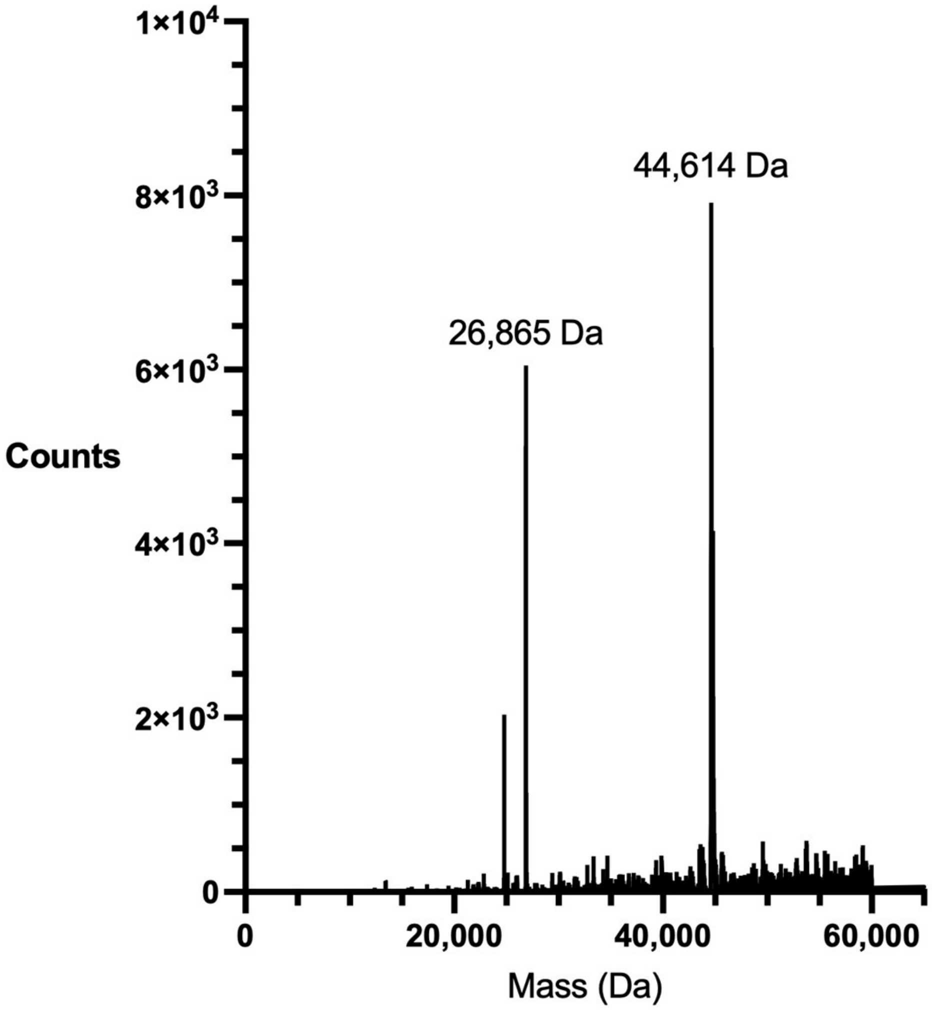


***Supplementary Figure 2*. Intact mass spectrometry of NanoBondy**.
Electrospray ionization mass spectrometry of anti-CD45 2H5 R72C NanoBondy. Expected mass of the full-length NanoBondy after disulfide formation is 44,614 Da. Expected mass of the SPM cleavage product is 26,865 Da.

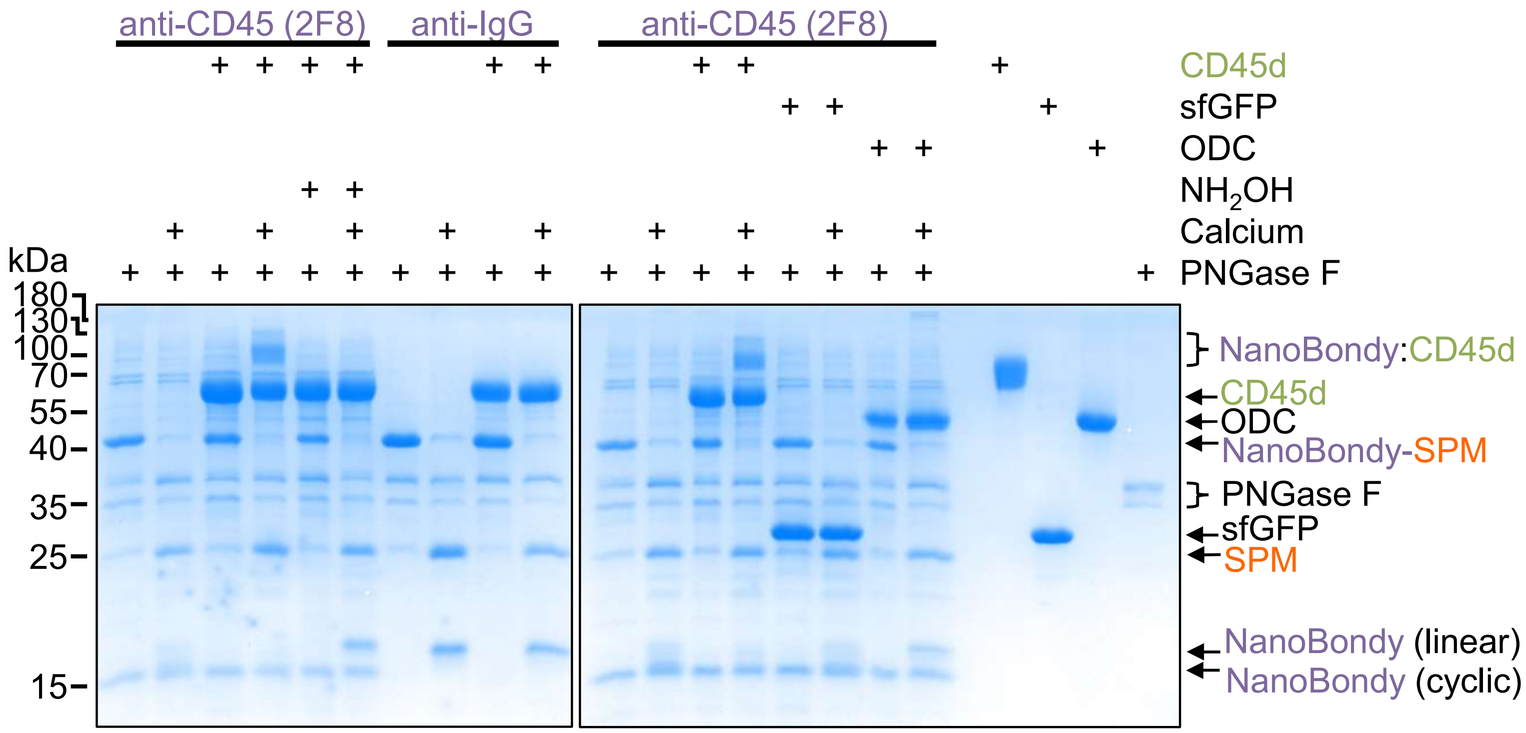


***Supplementary Figure 3.*** **Specificity of 2F8-derived NanoBondy coupling to CD45**. α-CD45 2F8 R72C NanoBondy was incubated ± calcium with CD45d or control proteins ornithine decarboxylase (ODC) or superfolder GFP (sfGFP) for 1 h at 37 ºC in HBS. Hydroxylamine was a competitor. α-IgG NanoBondy was a negative control. Formation of the covalent product was visualized using SDS-PAGE/Coomassie.

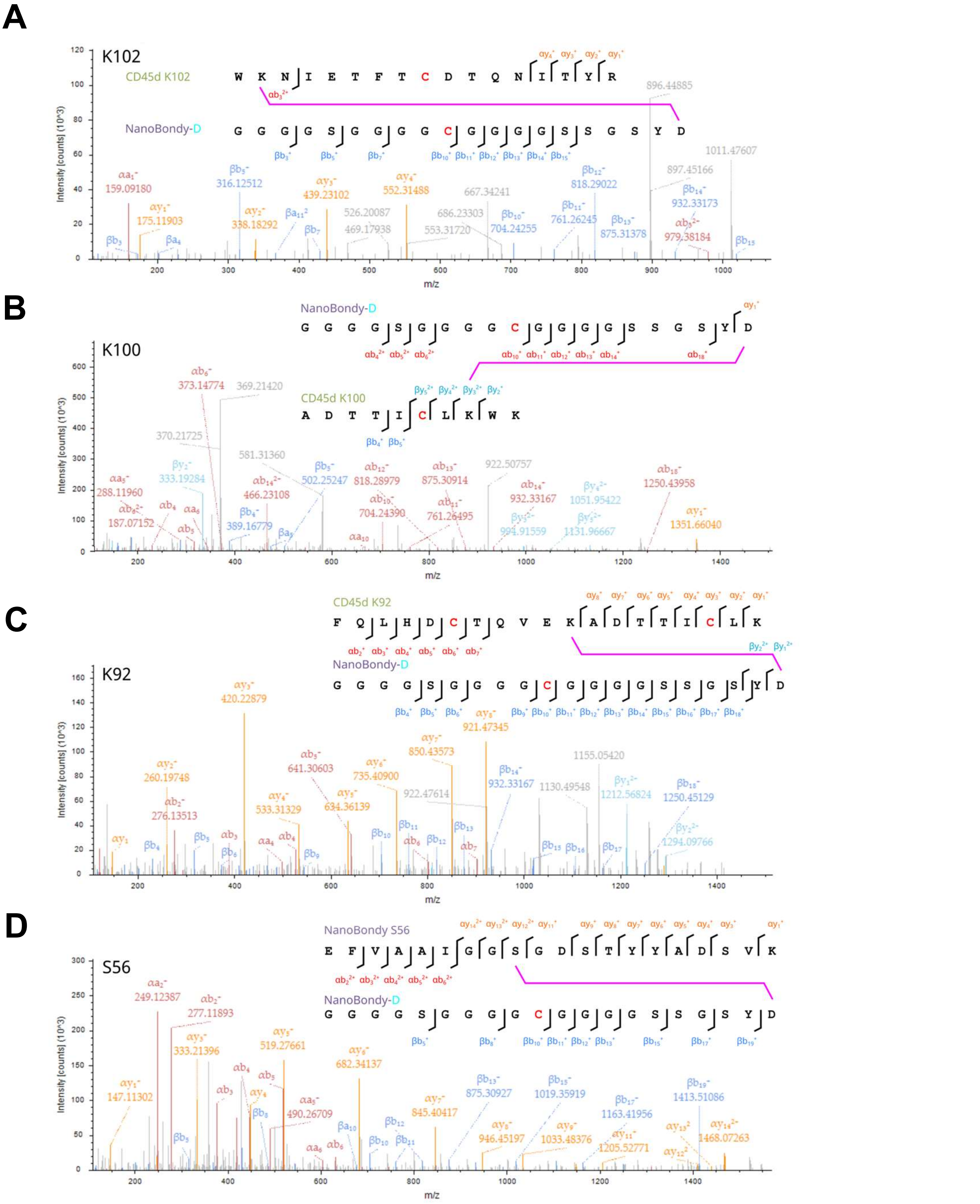


***Supplementary Figure 4.*** **NanoBondy crosslinks mapped in MS/MS.** Higher energy collision-induced dissociation (HCD) fragmentation spectra of identified crosslink precursor ions corresponding to D173 (2H5 R72C NanoBondy) peptide coupled to a CD45d peptide: **(A)** K102, **(B)** K100, **(C)** K92 and self-link corresponding to NanoBondy D173 coupled to **(D)** NanoBondy S56. MS/MS spectra taken directly from Proteome Discoverer. Cysteines are colored red to indicate carbamidomethylation. The two peptides are depicted as α and β peptides (α peptide above β peptide in figure). α-b (red), α-y (orange), β-b (dark blue) and β-y (light blue) ions are marked. Some spectra also contain a ions, which were ignored for fragment annotation.

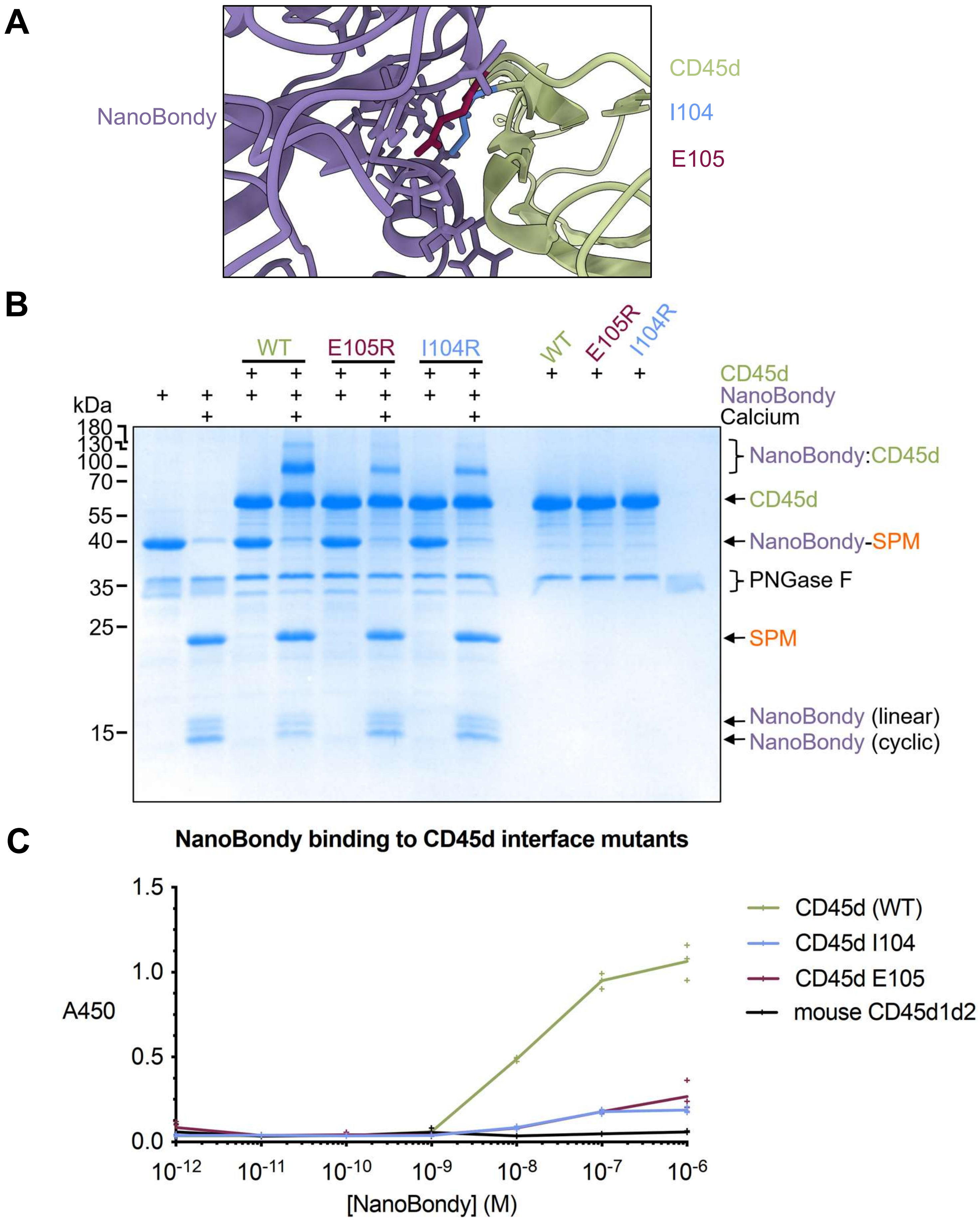


***Supplementary Figure 5.*** **Validation of predicted NanoBondy docking site on CD45. (A)** AlphaFold2multimer prediction of 2H5 NanoBondy:CD45d docking. E105 (teal) and I104 (blue) of CD45 sit at the docking interface. NanoBondy residues 33-36 and 103-107 shown in stick form. **(B)** 2H5 R72C NanoBondy conjugation to CD45d mutants was evaluated by SDS-PAGE/Coomassie. 10.5 µM of NanoBondy was coupled with 10.5 µM of CD45d variants for 1 h at 37 ºC ± $Ca^{2+}$, then digested with PNGase F for 2 h. PNGase F was included in all lanes. **(C)** NanoBondy binding to CD45d mutants evaluated using ELISA. WT CD45d, CD45d mutants, or mouse CD45d (mCD45d1d2, negative control) were bound to the plate and incubated with a titration of NanoBondy variants. NanoBondy binding was detected using anti-VHH HRP. Each triplicate data point is shown, with a line connecting the mean.
